# Sephin1 rewires proteostasis through actin-dependent signaling

**DOI:** 10.64898/2026.04.20.719601

**Authors:** Giulia Frapporti, Antonella Capuozzo, Eleonora Colombo, Paolo Fioretti, Vincenzo Maria D’Amore, Francesco Saverio Di Leva, Adriano Lama, Vasvi Tripathi, Stefano Medaglia, Stephanie Waich, Caterina Montani, Maria Dolores Perez-Carrion, Antonella Marte, Franco Onofri, Christian Johannes Gloeckner, Luciana Marinelli, Pierfausto Seneci, Michael W. Hess, Diego L. Medina, Giovanni Piccoli

**Author notes:** equal contribution (Michael W. Hess, Diego L. Medina).

## Abstract

The maintenance of protein homeostasis is vital for all cells. Alteration in protein handling underlies several diseases. The small molecule sephin1 is a promising clinical candidate against proteostasis disruption, but its mechanism of action is still uncertain. Our experimental evidence shows that sephin1 binds G-actin and drives actin cytoskeleton misfolding, and eventually, Golgi disintegration. At first, sephin1 impairs the autophagic flux and elicits the phosphorylation of the α subunit of eIF2 and the ER-stress independent expression of CHOP via GCN2 kinase. Sephin1 also inhibits the mammalian target of rapamycin (mTORC1), activates the transcription Factor EB (TFEB), drives the expression of TFEB-direct target genes, and eventually stimulates the autophagy lysosomal pathway. Our results reveal that the actin cytoskeleton may regulate autophagy via mTORC1-TFEB complemented with the GCN2-eIF2α-CHOP signaling pathway.

## Introduction

The cell utilizes an extensive and complex network of signalling pathways to preserve its proteome’s homeostasis, the proteostasis network (PN). The PN guarantees that superfluous and misfolded protein species are removed: PN unbalance affects cellular protein quality control and leads to accumulation of toxic protein aggregates, and, eventually, age-related neurodegenerative diseases ^1–3^. Translation attenuation is critical in relieving PN overload in conditions of conformational stress, such as when misfolded or unfolded proteins accumulate in the ER lumen. Generally, translation attenuation is mediated by the inhibition via phosphorylation of the translation initiation factor 2α (eIF2α) via different kinases, including PERK and GCN2. The phosphorylation of eIF2α (p-eIF2α) marks the beginning of a cytoprotective pathway called the integrated stress response (ISR). Under diverse stressful conditions, the α-subunit eIF2 is phosphorylated on Ser52 (or Ser51 in rodents) in its N-terminal domain. Generally, eIF2α is the substrate of eIF2B-guanine-nucleotide-exchange (GEF)-activity but once phosphorylated is converted to an inhibitor of eIF2B. eIF2B mediates the exchange of eIF2α-GDP to eIF2α-GTP, thus promoting the binding of initiator methionyl-tRNA (Met-tRNAi) to eIF2α-GTP and, eventually, enhancing protein translation ^4^. By depleting the eIF2α/GTP/Met-tRNAi ternary complex, eIF2α phosphorylation induces the reduction of global protein synthesis. At the same time, eIF2a phosphorylation promotes the selective expression of genes via non canonical re-initiation mechanisms ^5^. One key target is the activating transcription factor 4 (ATF4) ^6^. ATF4 promotes the expression of the C/EBP homologous protein (CHOP). ATF4 and its target CHOP, alone or even in combination, guide the transcriptional upregulation of several genes involved in protein stress resolution response, including the autophagy lysosomal pathway (ALP) ^7,8^. ALP is regulated by a wide range of cellular signals, such as cellular stress, nutrient starvation, growth factors, hypoxia, or ER stress. The first sensor of amino acid, growth factor and hormone variation is the mammalian target of Rapamycin (mTOR), an evolutionary conserved serine/threonine protein kinase first described by Heitman ^9^. To be active, mTORC1 must localize at the lysosomal membrane, where its co-activator RHEB-GTP resides. In nutrient-rich conditions amino acids are sensed at the lysosome membrane by the vacuolar-type H+-translocating ATPase in conjunction with RRAG proteins and the Ragulator complex; together they direct mTORC1 to the lysosome membrane where it becomes activated inhibiting ALP induction ^10,11^. Conversely, under conditions of amino acid withdrawal, mTORC1 is inhibited and ALP starts. The major well-characterized mTORC1 substrates include eukaryotic translation initiation factor 4E (eIF4E)-binding protein 1 (4E-BP1), ribosomal protein S6 kinase 1 [S6K1, also known as p70S6 kinase (p70S6K, p70-S6K)], ULK1, Atg13, and TFEB ^12–14^. TFEB is among the master regulators of the ALP. Unphosphorylated TFEB translocates to the nucleus where it promotes the transcription of a large subset of lysosomal and autophagic genes ^15^. Overall, protein stress leads to the sequential activation mechanisms involving attenuation of global protein synthesis, transcriptional induction of genes, and ultimately leads to autophagy induction ^16^. Such coordinated response holds crucial therapeutic implications. Strategies to target specific components of the PN using small molecules and gene therapy have been developed and represent interesting avenues for future interventions to delay or eventually stop proteinopathies, including neurodegenerative diseases. The phosphorylation of eIF2α is an evolutionarily conserved defense mechanism against protein stress. Consequently, eIF2α phosphatase inhibitors have been tested for their therapeutic potential.

Guanabenz (GBZ) and its analogue sephin1 (*IFB-088/Icerguastat*) were postulated to bind PPP1R15A, thus inhibiting p-eIF2α dephosphorylation and enhancing ISR signaling ^17,18^. Sephin1 shows therapeutic effect in several CNS disease mouse models of ALS, multiple sclerosis, prion disease, Charcot-Marie-Tooth disease, and ischemia ^19–23^. Sephin1 has entered clinical trials as a standalone orphan drug ongoing Phase II study and in combination with riluzole on patients with bulbar-onset ALS [https://clinicaltrials.gov/study/NCT05508074]. However, subsequent studies failed to confirm either GBZ and sephin1 binding to PPP1R15A or impairment of p-eIF2α dephosphorylation ^24,25^. Thus, there is an unmet need to identify the true molecular interactor for the rational development of next-generation proteostasis modulators. Here, we pivot from the phosphatase-centric model to propose monomeric actin as the primary target of sephin. Acutely, sephin1 causes the remodelling of the perinuclear actin cytoskeleton via Arp2/3, Golgi fragmentation, and a transient block of autophagy. The initial actin remodelling start a signalling cascades encompassing both the eIF2-CHOP and mTOR-TFEB pathways. Eventually, sephin1 induces an overt stimulation of the ALP. Together, we provide a mechanistic bridge between cytoskeletal integrity and the integrated stress response (ISR) upon sephin1 treatment.

## Material and methods

### Cell lines, transfections and drugs

HeLa cells (ATCC: CCL-2) were cultured in DMEM (Euroclone) medium, supplemented with 10% fetal bovine serum (FBS; Euroclone), 1% penicillin/streptomycin (Gibco, 15140122) and 1% L-glutamine (Gibco) in a humidified atmosphere of 5% CO2 at 37°C. Cells were passaged one or two times a week depending on confluency using Trypsin-EDTA 0.05% (Gibco, 25300054). HeLa TFEB-GFP cells were described in (PMID: 21617040). Cells were silenced using 40 nM of siRNAs against TRPML1 (sequences 5’-3’: #1 CCUUCGCCGUCGUCUCAAA; #2 AUCCGAUGGUGGUUACUGA; #3 GAUCACGUUUGACAACAAA), and against Calcineurin, catalytic subunit PPP3CB (Ambion, 4390824), and regulatory subunit PPP3R1 (Invitrogen, HSS108410-3, HSS183046-3, HSS108411-3) for 72 hours using the Lipofectamine RNAimax (ThermoFisher) reagent accordingly to the protocol provided by the manufacturers. PPP1R15B-Flag (a kind gift from David Ron laboratory), LifeAct-mCherry (Addgene, #193300), GFP-Golgi (Addgene, #182877), GFP-LC3 (Addgene, #11546constructs) were transiently transfected with Jet-PEI (Polyethylenimine, Polysciences) with a 0.8:100 DNA:PEI ratio. Cultures were treated at 37 °C for the time indicated with the drugs listed in Table 1.

**Table 1.**
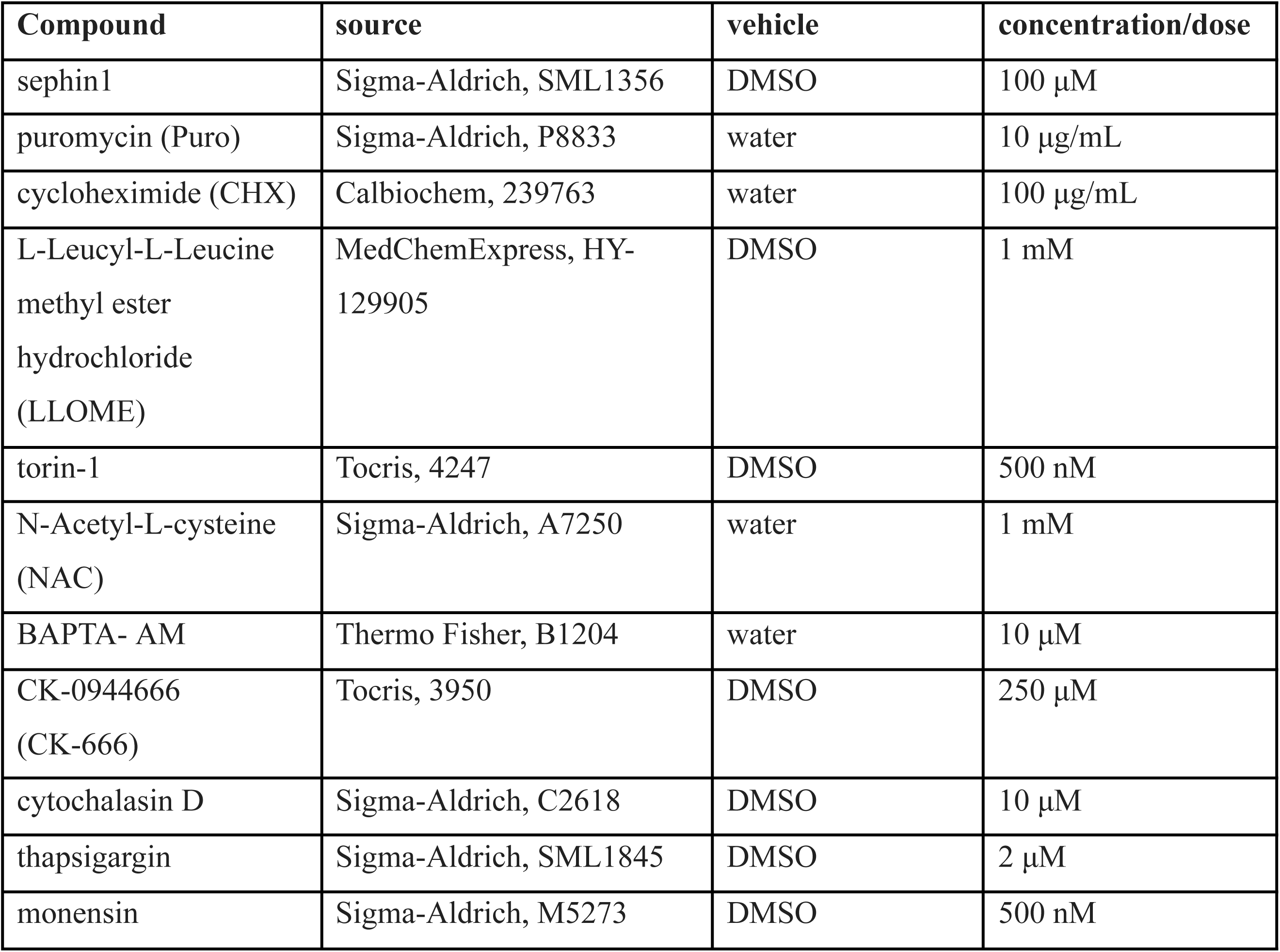
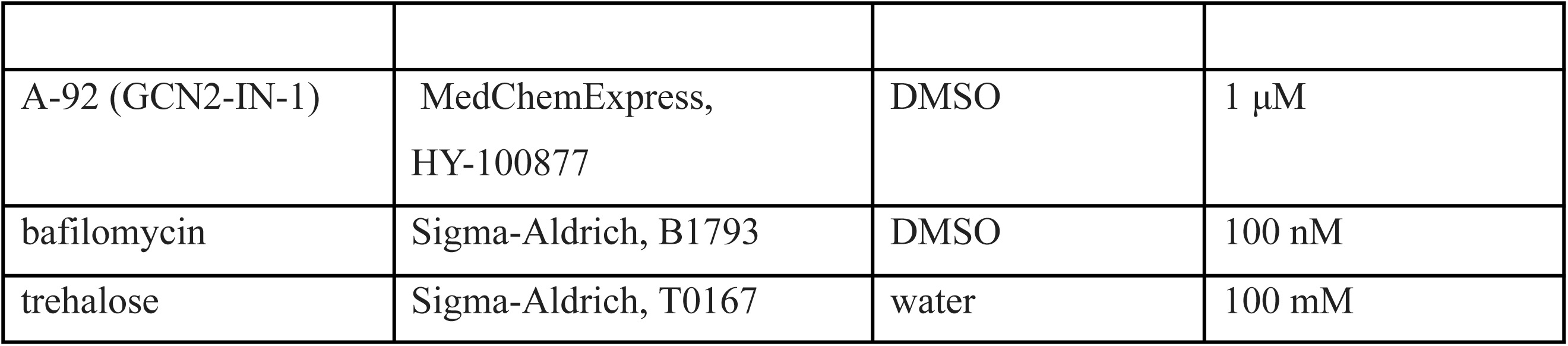
List of drugs.

### Cytotoxicity assay

We performed the 3-(4, 5-dimethylthiazol-2-yl)-2,5-diphenyltetrazoliumbromide (MTT) assay to measure cell vitality. HeLa cells were cultured in a 96-well plate at a concentration of 5 x 10^3^ cell/cm^2^ and incubated at 37 °C for 2 hours. Afterwards, cell medium was removed and MTT was added at a final concentration of 0.25 mg/mL in PBS. After an incubation of 30 minutes at 37°C, the MTT solution was carefully aspirated, and formazan precipitates were collected in 200 μL of DMSO. The absorbance measured at 570 nm using a spectrophotometer reflected cell viability, expressed as fold over control condition set at 1.

### RNA extraction and quantitative real-time PCR

Total RNA was extracted from cells using the Total RNA Purification Kit (Norgen), according to the manufacturer’s protocol. After extraction, RNA concentration was determined with the NanoDrop 2000C spectrophotometer (Thermo Fisher Scientific). Reverse transcription was performed using qRT SuperMix (Bimake) and following the instructions of the manufacturer. Real-time quantitative Reverse Transcription PCR (qPCR) was accomplished using iTaq Universal SYBR® Green Supermix and CFX96 RealTime System (BioRad), according to the BioRad recommendations. HPRT, GAPDH and rRNA18S were used as reference genes. Data was produced using Bio-Rad CFX Manager software. Expression levels data (Ct values) were analysed as follows. Each sample was run in triplicates, the geometric mean calculated and used for further calculations. To quantify the differences in gene expression we applied the Livak Method ^26^. Primers are listed in Table 2.

**Table 2.**
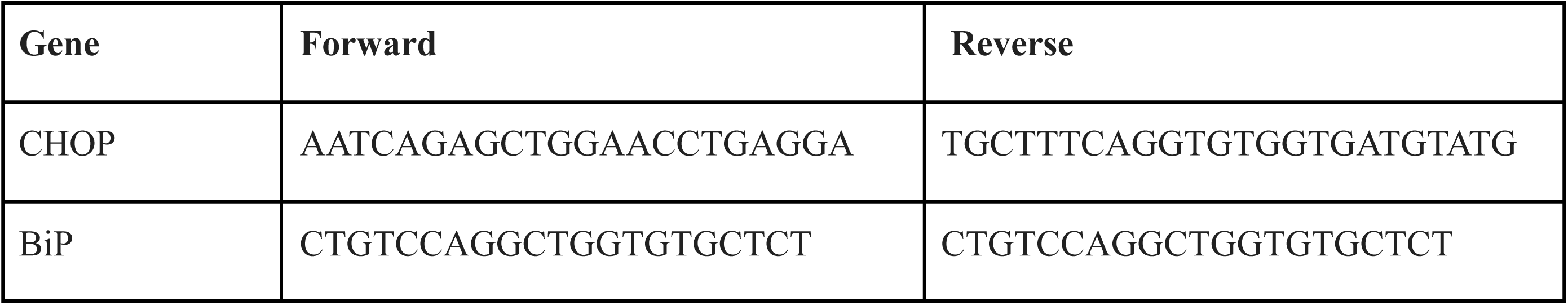
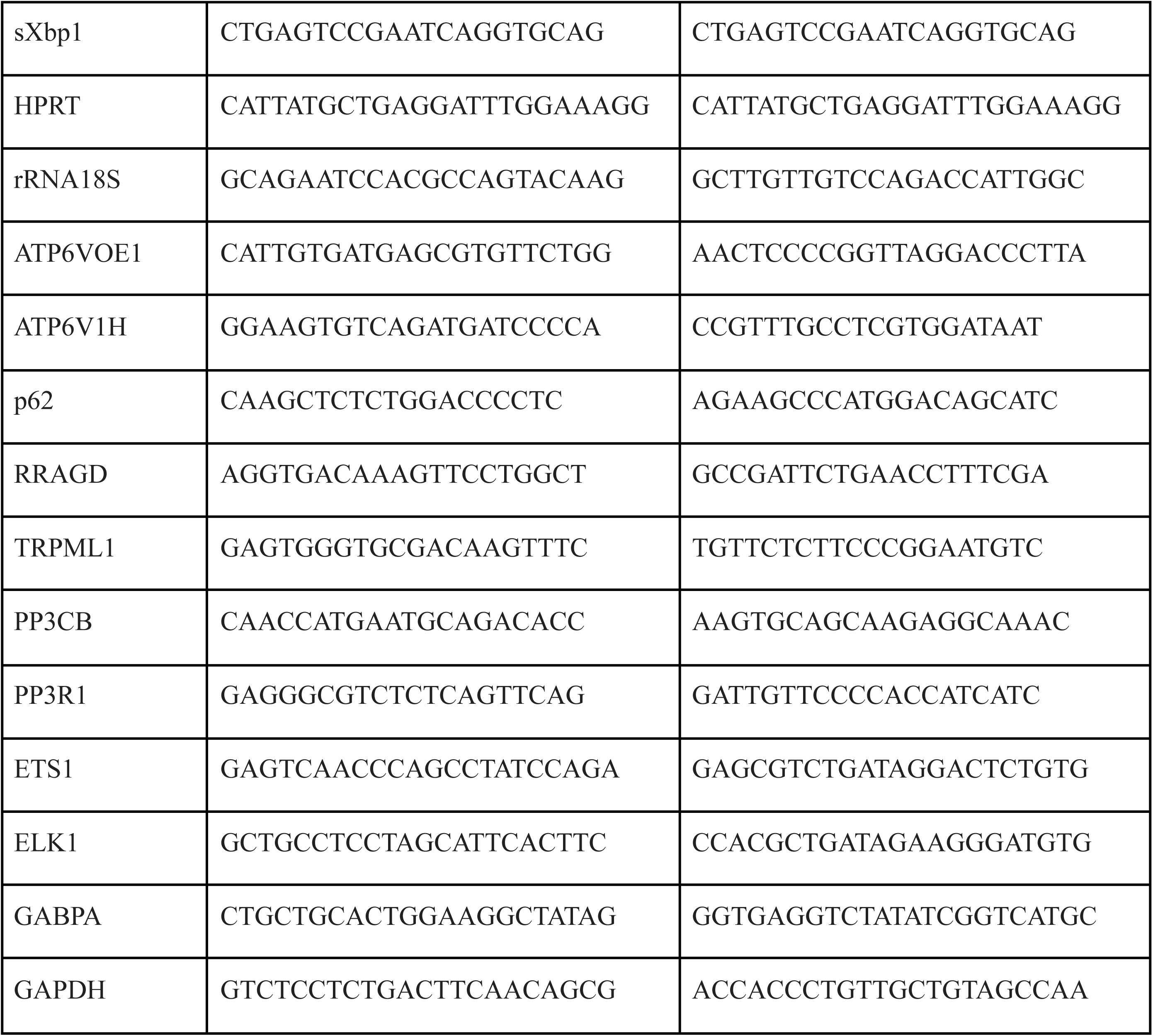
List of primers (5’-3’)

### Immunofluorescence

HeLa cells were seeded at low density on ø12mm Cover Glasses (Thermo Fisher Scientific) and fixed in 4% PFA. For the DQ-BSA assay, the cells were previously incubated overnight with 10 μg/mL of DQTM Red BSA added in normal medium. Prior to labelling, cells were permeabilized for 10 minutes in (2.5% BSA, 10% goat serum, 0.2% Triton-X100 and 0.05% saponin. Then, cells were incubated for 1 hour with either Alexa FluorTM 568 Phalloidin dye (1:400, Thermo Fisher, #A12380) or anti GAL3 antibody (1:400. Santa Cruz, SC-23938). After three washes of 10 minutes each with PBS, cells were incubated with fluorescent secondary antibodies in a dedicated buffer (2.5% BSA in PBS, 0.2% Triton-X100) for 45 minutes and then washed three times with PBS. Nuclei were stained with DAPI (NucBlue^TM^ Fixed Cell ReadyProbes^TM^ Reagent, ThermoFisher, R37606). Slides were mounted using FluorSave^TM^ Reagent (Millipore). For the analysis, at least 10 images were acquired per well by using the Opera system (Revvity) with a 40X objective. Images were analysed with ImageJ Software. To appreciate Golgi fragmentation, we divided the total perimeter of GFP-Golgi structures by the perimeter of each GFP-Golgi positive objects within each cell as in ^27^. The quantification of spots per cell was performed by Signals Image Artist (SImA)^TM^ Image Data Storage and Analysis System (Revvity). To access GFP-TFEB translocation, a dedicated script was developed to quantify the ratio derived from the mean intensity of GFP signal in a fixed ring region around the perinuclear area and the nucleus.

### Electron microscopy

HeLa cells were cultured for 3 days on carbon-coated sapphire discs and optionally treated with Sephin1, BafA1 and/or CK666 as indicated. Cells were eventually subjected to rapid cryofixation through means of high-pressure freezing, freeze-substitution and epoxy resin embedding ^28,29^. We performed immunoelectron microscopy with anti-actin (MAB1501R, 1:50), or anti-LC3 (PM036, 1:200, cosmobio##1:100) by using two different approaches, namely pre-embedding immunogold labelling of the cryofixed samples ^28^ and the Tokuyasu’s thawed cryosection labelling of formaldehyde-fixed samples ^30^. Pre-embedding labelling of cryofixed samples was essentially performed as previously described with minor modifications ^28^. Instead of Nanogold™ conjugates we used Alexa-488™ conjugated secondary antibodies (Invitrogen goat anti-rabbit, anti-mouse; A-11008, A-1100; 1:50), that were visualized after resin embedding by labelling the 100nm-thin sections with rabbit anti-Alexa 488 (1:50), followed by anti-rabbit 5 or 10nm colloidal gold conjugates (EM GAR5, GAR10, British Biocell Intl., 1:50). For Tokuyasu-labelling cells grown in petri dishes were fixed with 4% (w/v) buffered formaldehyde solution and bound primary antibodies were visualized with Nanogold™-conjugated secondary antibodies (NANOGOLD-Fab′ goat anti-mouse/-rabbit/ IgG (H + L), followed by silver enhancement (HQ-Silver, Nanoprobe). The reliability of the primary antibodies in immunoelectron microscopy approaches was validated previously ^28,31^. Labelling controls included among others cell samples treated with BafilomycinA in amino acid- and serum-free culture medium. Ultrathin sections (≈100nm thick) were analyzed with a CM120 transmission electron microscope (Philips, Eindhoven, The Netherlands) equipped with a MORADA G1 digital camera (EMSIS, Münster, Germany). Digital electron micrographs taken with iTEM-FIVE software (EMSIS) were optionally processed with Adobe Photoshop V.9, namely adjustment of contrast and brightness, gray-scale modification, sharpening, median and high-pass filtering.

### Pull-down assays

HeLa cells were lysed in RIPA buffer (150mM NaCl, 50mM HEPES, 0,5% NP40, 1% Sodium-deoxycholate) for one hour at 4°C and then processed for streptavidin immunoprecipitation in the presence of biotin or FG08 (10 and 100 μM). Human platelet non-muscle actin (Cytoskeleton Inc., APHL99-C) was diluted to a final concentration of 250 nM in the actin buffer [5 mM Tris-HCl pH 8.0, 0.2 mM CaCl2, 0.2 mM ATP, 5% (w/v) sucrose, and 1% (w/v) dextran]. We incubated diluted actin on ice for 60 minutes and possible aggregates formed during storage were cleared via ultracentrifugation (100 000 x g for 1 h at 4°C). The supernatant was transferred to clean microfuge tubes. FG08 and sephin1 were incubated with 250nM actin overnight at 4°C under gentle rotation. The next day, samples were incubated with Strep-Tactin Superflow resin (Iba-Lifescience) for 2 hours at 4°C. The resin was then extensively washed with actin buffer. Interacting proteins were eluted in Laemmli buffer 2X at 95°C for 10 minutes and subjected to 10% SDS-PAGE followed by silver staining or western-blotting.

### In vitro precipitation assay

Human platelet non-muscle actin (Cytoskeleton Inc., APHL99-C), rabbit skeletal muscle actin (Cytoskeleton Inc., # AKL95) or BSA (Sigma, A9647) were diluted to the concentration indicated in the text in the following actin buffer [2.5 mM Tris-HCl pH 8.0, 0.1 mM CaCl2, 0.1 mM ATP, 2.5% (w/v) sucrose, 0.5% (w/v) dextran]. Diluted actin was incubated on ice for 60 minutes and cleared by ultracentrifugation (100 000 x g for 1 h at 4°C). The supernatant was transferred to clean microfuge tubes. Target proteins were incubated with sephin1 with or without 0.01 % Triton-X100. Upon centrifugation at 16 000 x g for 15 minutes at 4°C, the supernatant and pellet were collected and analysed by western blotting.

### Western blotting

Total cell lysates were prepared by solubilization of samples in RIPA buffer (150 mM NaCl, 50 mM HEPES, 0.5% NP40, 1% sodium-deoxycholate) supplemented with protease inhibitors (Calbiochem) and phosphatase inhibitor cocktail (Sigma Aldrich). The samples were briefly vortexed three times every 10 minutes and then clarified by centrifugation at 10 000xg for 10 minutes. All experimental procedures were performed at 4° C. Protein concentrations of the lysates were determined via Bradford assay (Bio-Rad, Segrate, Italy). Either samples, blank control, or standards (BSA 1μg/ μL) were added to 1 mL Bradford reagent containing cuvette for protein concentration measurement. Absorbance was measured using Tecan plate reader (Tecan M200) at 600 nm and protein concentration was determined using a linear regression of the standard curve. For western blotting experiments, an equal amount of proteins was diluted with 5X Laemmli buffer, heated at 95°C for 10 minutes and loaded onto SDS-PAGE gels. After gel electrophoresis, the proteins were transferred on PVDF membranes. The transfer at 4°C lasted 90 minutes at 300 mA. Transfer efficiency was controlled with Ponceau S staining, then membranes were washed in ddH2O and finally in 1x TBS (20 mM Tris pH 7,4, 150 mM NaCl), before they were blocked in 5% non-fat dried milk in 1x TBS-Tween 1% (TBS-T) for one hour. Primary antibodies were incubated in blocking buffer overnight at 4°C on an orbital shaker. Primary antibodies used are listed in Table 3. After three washing steps in TBS-T, the secondary antibodies (HRP-conjugated anti-mouse, Jackson ImmunoResearch, UK; HRP-conjugated anti-rabbit, 1:5000) were applied for 60 minutes at room temperature. Then the membrane was washed three times in TBS-T. Proteins were detected using the ECL detection system (GE Healthcare) and chemiluminescence images were acquired using ChemiDoc Touch system (Bio Rad Laboratory Italy, Segrate, Italy). Optical density of the specific bands was measured with ImageLab software (Bio Rad).

**Table 3.**
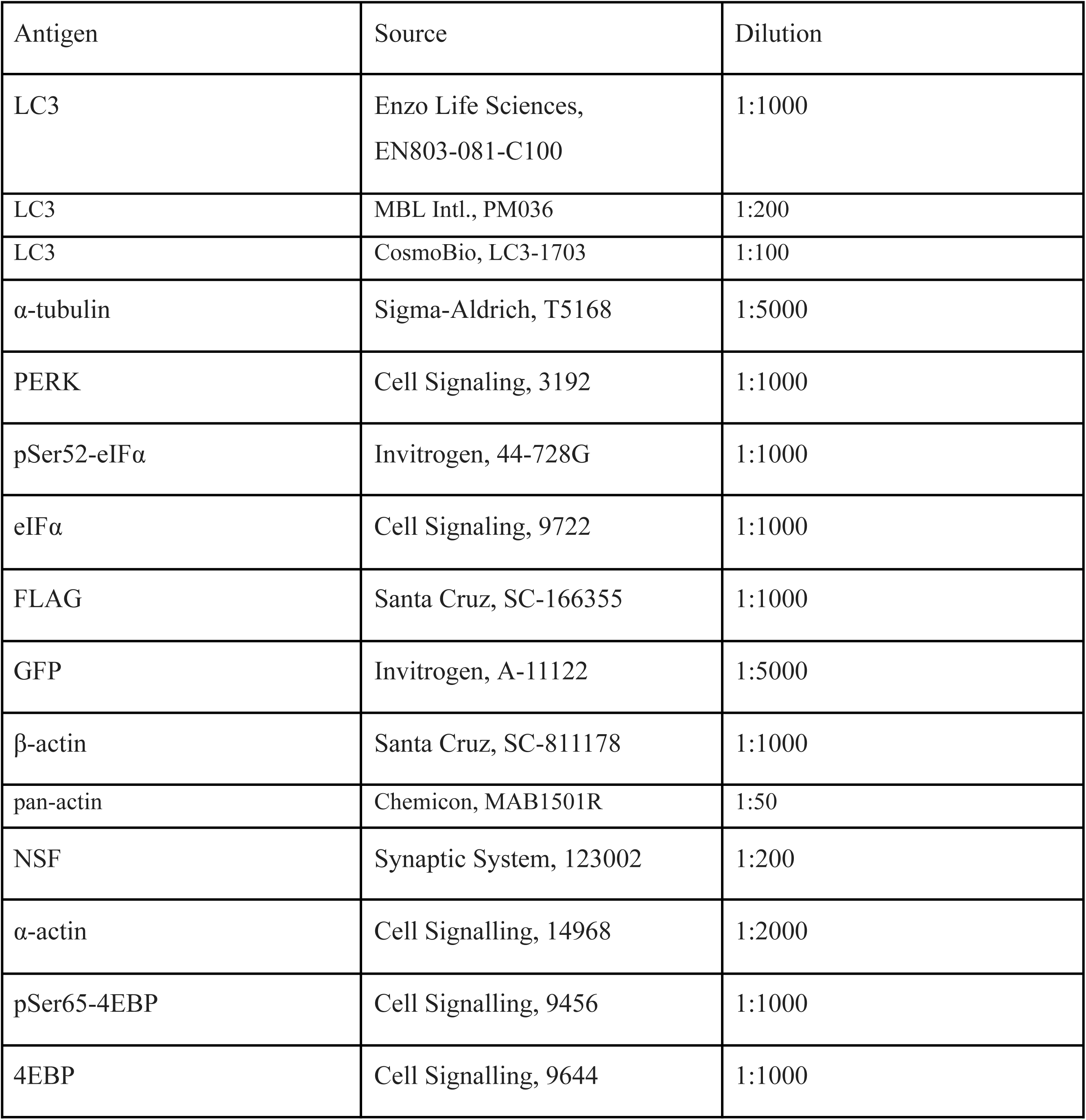
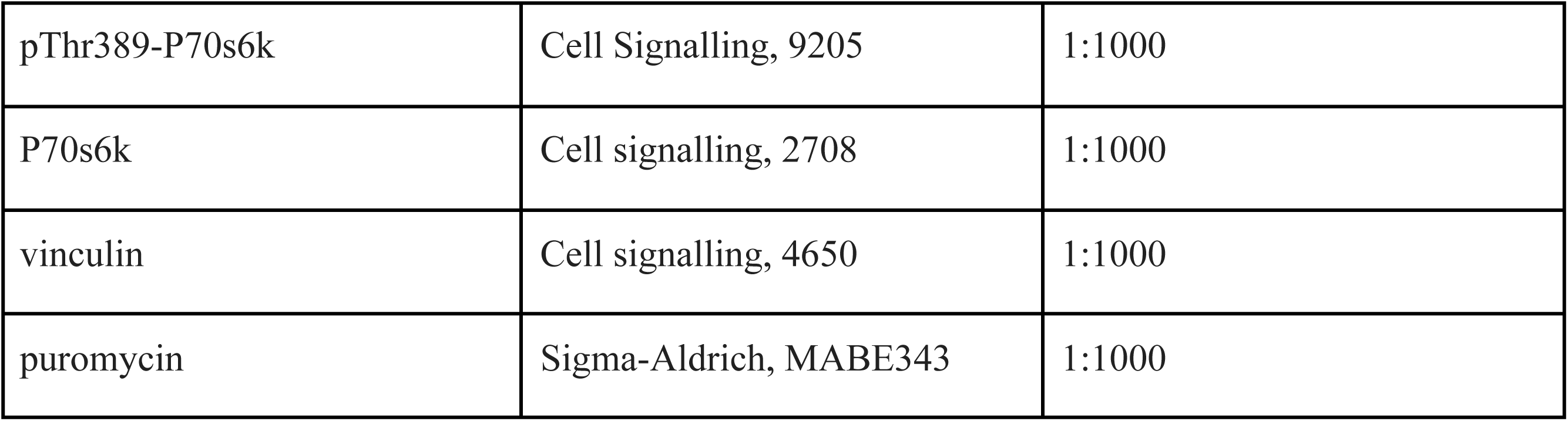
List of primary antibodies.

### Mass Spectrometry analysis

After Silver or Coomassie staining, the protein band of interest was cut from the gel. The excised gel band was cut into small pieces and subjected to reduction and alkylation with 10mM dithiothreitol (DTT) and 55mM iodoacetamide (IAA), respectively. The gel pieces were dried in a speed vacuum and incubated overnight at 37°C in 50 μL of digestion buffer containing 12.5 ng/μL of Trypsin (Thermo Scientific). The resulting peptides were extracted sequentially from the gel slices with 30% acetonitrile (ACN)/ 3% trifluoroacetic acid (TFA) and 100% ACN. All supernatants were combined and dried in a SpeedVac. Peptides were then acidified with 1% TFA, desalted on C18 stage-tips and resuspended in 15μL of 0.1% formic acid buffer for LC-MS/MS analysis. In-gel digested samples were analyzed using an Easy-nLC 1200 system, coupled online with an Orbitrap Fusion Tribrid mass spectrometer (both Thermo Fisher Scientific, San Jose, CA, USA) in data-dependent mode (DDA). A reversed-phase column (Thermo Fisher Scientific, Acclaim PepMap RSLC C18 column, 2μm particle size, 100Å pore size, id 75 μm) and a two-component mobile phase system of 0.1% formic acid in water (buffer A) and 0.1% formic acid in acetonitrile (buffer B) was used for separating the digested peptides. Peptides were eluted using a gradient of 5% to 25% over 52 minutes, followed by 25% to 40% over 8 minutes and 40% to 98% over 10 minutes at a flow rate of 400 nL/min. Ion source parameters were as follows: spray voltage +2500V, sweep gas 0 Arb, ion transfer tube temperature 275°C. The DDA method was based on full scan acquisitions performed in the Orbitrap at 120 000 FWHM resolving power (at 200 m/z). The AGC target is set at 1e6 and a maximum injection time of 50 ms. A mass range of 250-2 000 m/z was surveyed for precursors, with the first mass set at 140 m/z for fragments. Full scans were followed by a set of (HCD) MS/MS scans over 3 sec cycle time, at collision energy of 27% using the Ion Trap mass analyzer (35 ms of maximum injection time, AGC 1e4) or the Orbitrap (30 000 fwhm, 54 ms of maximum injection time, AGC 5e4). The dynamic exclusion filter was set at 20 sec, and the intensity of precursors for fragmentation filtered at 5e3. Data were acquired using Xcalibur 4.3 software and Tune 3.3 (Thermo Scientific). For all acquisitions, QCloud was used to control instrument longitudinal performance during the project using in-house quality control standards ^32^. Raw data were searched in ProteomeDiscoverer 2.2TM (Thermo Scientific). Peptides were searched using MASCOT algorithm against the in-silico digested P60709 and P63261 proteins sequence (uniprot, downloaded April 2021). A 10 ppm and 0.6 Da mass tolerance values were selected for precursors and products, respectively. Trypsin digestion was performed, with up to 5 missed cleavages. Fixed modification of carbamidomethylation of cysteine residue (+57.021Da) and variable modifications of oxidation of methionine (+15.995Da) and acetylation of protein N-terminus (+42.011Da) were incorporated in the search, as well as the addition of the MW of the irradiated PAL derivative EC186 (C_22_ClF_3_H_20_N_8_O_2_, corresponding to 520.134984 monoisotopic mass shift, see supplementary figure 5A). Peptide sequences were searched if their lengths were longer than or equal to 6 aa.

### Molecular Modeling

To investigate the interaction between sephin-1 and actin at the molecular level, we selected the cryo-EM structure of human monomeric ADP-bound β-actin (PDB code: 8COG [ https://doi.org/10.1083/jcb.202409067]) as the receptor model. The protein structure was prepared using the Protein Preparation Wizard ^33^ within the Maestro v.2025-2 software suite [Schrödinger, LLC, New York, NY, USA]. Missing hydrogen atoms were added, protonation states were assigned, and hydrogen-bonding networks were optimized at physiological pH. The positions of all hydrogen atoms were subsequently minimized using the OPLS-2005 force field^34^. The three-dimensional structure of sephin1 was generated using the 2D Builder module and prepared with LigPrep [Schrödinger, LLC]. Ligand protonation and tautomeric states within a pH range of 7.4 ± 1.5 were assigned using Epik ^35^. Docking calculations were performed using the grid-based Glide program ^36–38^. Prior to docking, potential druggable binding sites on β-actin were identified using the SiteMap algorithm ^39,40^. Among the predicted sites, the highest-ranked pocket corresponded to the ADP-binding site and was therefore excluded from further analysis. The second-ranked site was selected for docking because of its proximity to the peptide identified by PAL-LC–MS/MS analysis (Supplementary Fig. 5C). A receptor grid box (20 Å × 20 Å × 15 Å) centered on the selected binding site was generated using the Receptor Grid Generation tool. Docking simulations were performed in Standard Precision (SP) mode using the OPLS-2005 force field, while all other parameters were kept at their default values. The docking results showed clear convergence toward a predominant binding pose, as shown in Supplementary Fig. 5D.

### Statistical analysis

Data are expressed as the mean ± standard error of the mean (SEM). Statistical analyses were performed using GraphPad Prism. Unpaired two-tailed Student’s T-test was used when comparing two experimental groups. One-way ANOVA followed by Bonferroni’s multiple comparisons was used in experiments comparing three or more groups. Two-way ANOVA followed by Bonferroni’s multiple comparisons was used to analyse the influence of two different categorical independent variables. Normality distribution was always tested prior to applying a parametric test. For all analyses, the threshold for significance was set at p<0.05; * p<0.05, ** p<0.01, *** p < 0.001.

## Results

### Acute sephin1 treatment jeopardizes the autophagy flux

Complementary observations suggest that sephin1 influences the autophagic flux ^7,23^. Therefore, we evaluated the impact of sephin1 on autophagy by probing LC3B-I to -II conversion in presence of the lysosomal inhibitor bafilomycin A1 (BafA1). Based on pilot experiments we focussed first on for 2 hours incubations with 100μM Sephin1and/or 100nM BafA11 and measured LC3B accumulation. Sephin1 and BafA1 treatments yielded similar results, but the co-treatment with both drugs did not increase LC3B-II levels further (figure 1A-B). Sephin1 affects autophagy in a dose-dependent manner and is cytotoxic at concentration higher than 100 μM (supplementary figure 1A-F). Lysosomal damage or hypoactivity may cause an impairment in the autophagic flux. By probing galectin-3 staining to test lysosomal membrane integrity ^41^, we noticed that sephin1 does not trigger overt lysosomal damage (supplementary figure 1G-H). Electron microscopy (EM) of rapidly cryofixed cells revealed an accumulation of autophagic organelles upon sephin1 administration (figure 1D-I); predominantly a heterogenous population of (forming) autophagosomes (including cup-shaped phagophores/isolation membranes ^42,43^, but also amphisomes and autolysosomes, appear. We appreciated varying fragmentation and/or swelling of Golgi/TGN, but no ER dilatation. Lysosomes appeared structurally intact. By boosting eIF2 phosphorylation, sephin1 should reduce global protein synthesis but, at the same time, promote the expression of ISR proteins ^6^. However, the puromycin labelling assay did not record an over reduction of peptide elongation upon sephin1 treatment (supplementary figure 2A-B). Cycloheximide, a protein synthesis inhibitor, may affect autophagy ^44^. Cycloheximide (100 μg/ml, 2 hours) did not cause LC3-II accumulation or influenced sephin1 activity (supplementary figure 2C-H). Our results indicate that sephin1 impairs the maturation and clearance of autophagic organelles upon acute treatment in a protein synthesis independent manner.

**Figure 1.**
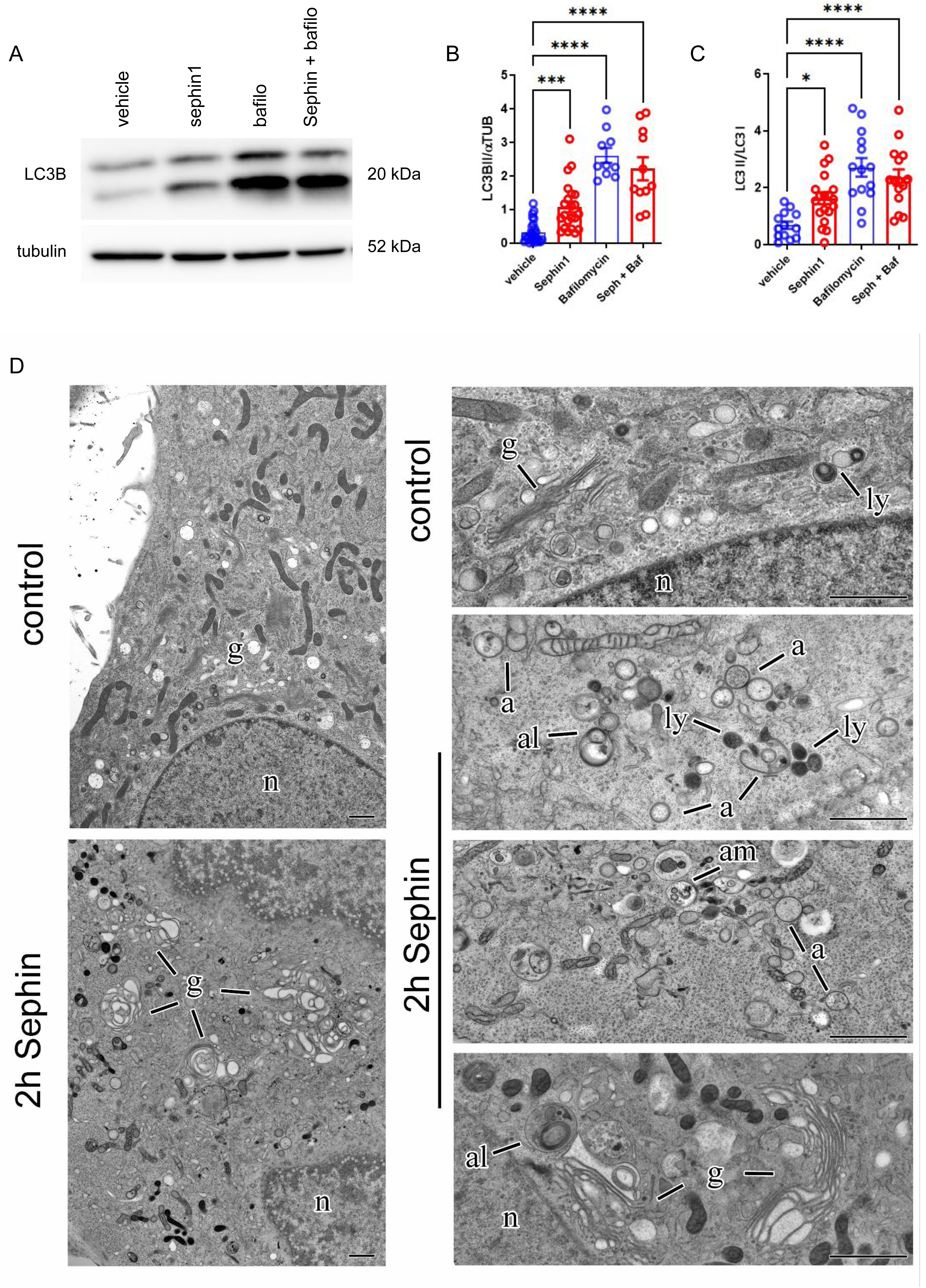
Sephin1 jeopardizes the autophagic flux. (A) We assessed autophagy by monitoring LC3I and LC3B II levels in HeLa cells treated for 2 hours with sephin1 (100μM) in the presence or absence of Bafilomycin 1 (BafA1, 100nM). (B-C) The graphs report the quantification of LC3B II levels, normalized to α-tubulin (B) and LC3II/LC3I ratio. Data are expressed as mean ± SEM, n =10-12. (D) (D) Electron micrographs of cryofixed HeLa cells at steady state (control) or upon Sephin 1 exposure (100μM, 2 hours). Sephin 1 treated cells appear with abundant (forming) autophagosomes (a), amphisomes (am) and autolysosomes (al) as well as vacuolized Golgi/TGN (g); (ly)=lysosomes, (n)=nucleus; scale bar= 1µm.

### Sephin1 induces Golgi stress

ER and Golgi play an important role in autophagy ^45–47^. CHOP, BiP, and sXbp1 mRNA levels monitor the presence of ER stress, and CHOP mRNA is upregulated upon eIF2a phosphorylation via ATF4. sephin1 caused an increase in CHOP mRNA levels but it did not alter the expression level of either BiP or sXbp1 (figure 2A). The observed activity of sephin1 has been originally explained by the inhibition of the stress-induced PPP1R15A and the consequent prolongation of eIF2α phosphorylation ^17^. We assessed the phosphorylation levels of eIF2α on Serine 52 (p-Ser52-eIF2α), PERK phosphorylation, and LC3B-I and II in HeLa cells, either in naive condition or upon treatment with sephin1 or the ER-stressor agent thapsigargin (2μM, 2 hours). Sephin1 treatment did not cause PERK phosphorylation but induced a significant increase in p-Ser52-eIF2α levels as well as LC3-II accumulation (figure 2B-F). Considering the altered Golgi morphology observed by EM, we next assessed sephin1 impact on the Golgi by biochemistry and fluorescence microscopy. Sephin1 treatment induced the expression of the Golgi stress markers ETS1 and GABPA (figure 2G) and caused Golgi disintegration (figure 2H-J) resembling the effects of the Golgi stressor monensin (500 nM, 2 hours) ^48^. Monensin promotes an autophagy-like process targeting damaged Golgi membrane (GOMED) in a bafilomycin sensitive manner ^49^. However, sephin1 activity on LC-3II was not influenced by pretreatment with bafilomycin (figure 2 K-M). Altogether, our data suggest that sephin1 causes Golgi stress and promotes LC3-II accumulation through an alternative mechanism.

**Figure 2.**
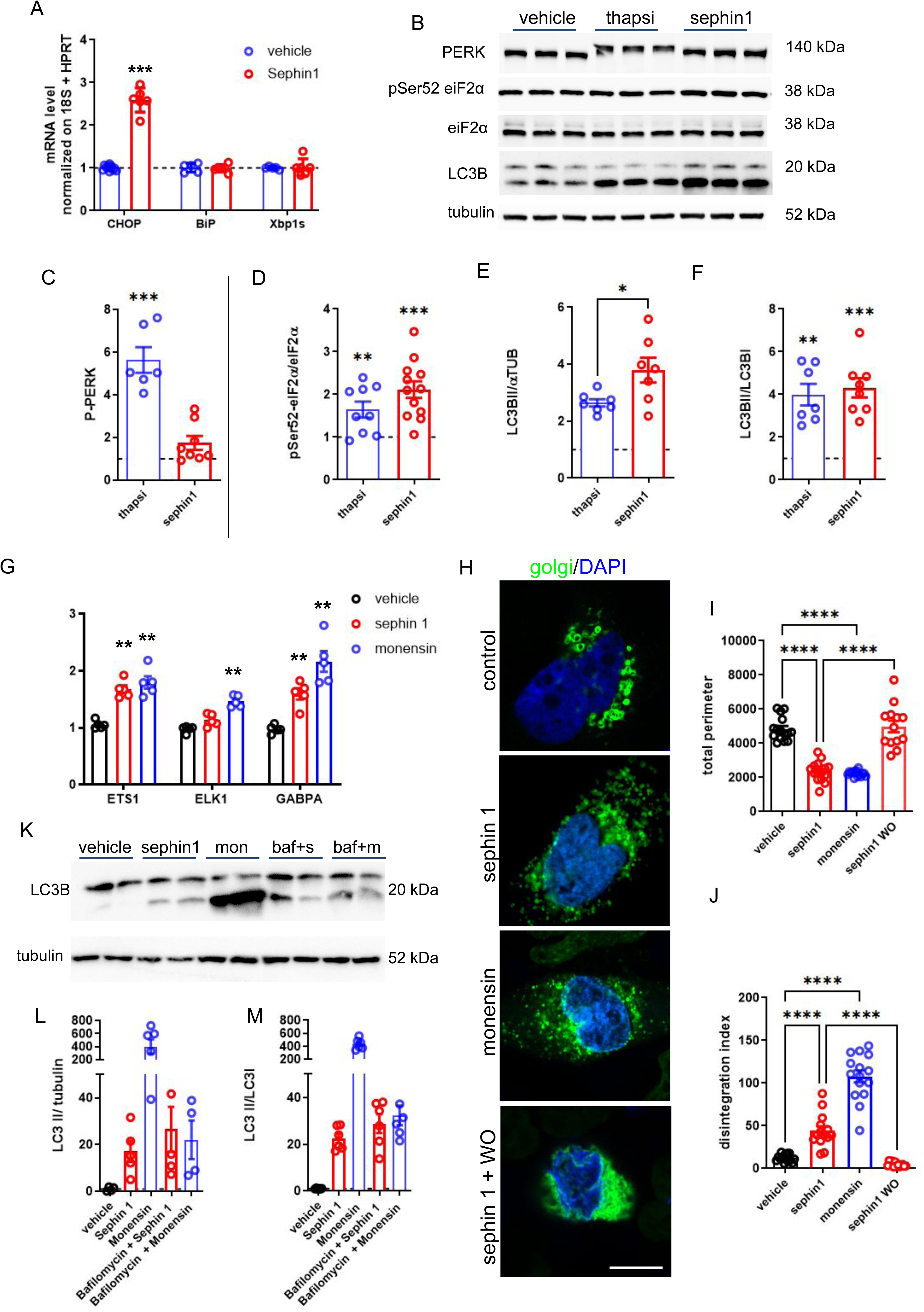
Sephin1 causes Golgi stress. (A) We profiled by qRT-PCR the expression of genes activated upon ER stress, such as CHOP, BIP, and Xbp1s upon sephin1 treatment (100μM, 2 hours). Data show 2^-△△Ct^ and are expressed as mean ± SEM, n =5. (B) We monitored by western-blotting PERK and eiF2α phosphorylation upon thapsigargin (thapsi, 500 nM, 2 hours) or sephin1 (100μM, 2 hours) treatment. (C-F) The graphs report the quantification of P-PERK (evaluated considering the electrophoretic shift) (C), eIF2a phosphorylation (D), LC3B II levels, normalized to α-tubulin (E) and LC3II/LC3I ratio (F). Data show fold over vehicle and are expressed as mean ± SEM, n =6-10. (G) We profiled by qRT-PCR the expression of genes activated upon Golgi stress, such as ETS1, ELK1, and GABPA upon 2 hours sephin1 or monensin treatment. (H) We monitored Golgi morphology by expressing the GFP-Golgi reporter in HeLa cells upon 2 hours of sephin1 or monensin treatment, scale bar 10 μm. (I-J) The graphs report Golgi total perimeter (I) and disintegration index (J). Data are expressed as mean ± SEM, n =10-12

### Sephin1 binds G-actin and alters its biochemical properties

Sephin1 has been described as a potent inhibitor of the PP1-PP1R15A complex (figure 3A). The over-expression of the PPP1R15B holoenzyme abolishes eIF2α phosphorylation ^50^. To study the role of the PP1c-PPP1R15A complex within autophagy, we evaluated LC3 conversion in cells overexpressing PPP1R15B or GFP as a control upon sephin1 treatment. Interestingly, we noticed that PPP1R15B overexpression did not buffer either LC-3II accumulation (figure 3B-E) or CHOP mRNA induction (figure 3F) observed upon sephin1 treatment. These data suggest that an alternative mechanism may link sephin1 to autophagy. Among the different kinases targeting eIF2 we focused on GCN2. The pharmacological inhibition of GCN2 impaired the eIF2 phosphorylation and CHOP mRNA induction triggered by sephin1 but did not overtly affect sephin1-induced autophagic impairment (figure 3G-K). To identify the sephin1 mechanism of action, we pull-down putative targets from HeLa protein lysate using biotinylated sephin1 as a bait (EC107, supplementary figure 3A). Upon immobilization on streptavidin resin, we characterized the eluted protein by SDS-PAGE followed by silver staining (supplementary Figure 3B). MS/MS analysis determined β-actin as putative interactor (supplementary Figure 3C). We confirmed this result by western blotting (figure 4A). To determine the nature of the sephin1-actin interaction, we incubated either purified monomeric β-actin or ɑ-actin (250 nM) with biotinylated-sephin1 (100 μM) in presence of 0.01% Triton-X 100. Upon streptavidin pull-down, we found that biotinylated-sephin1 binds to both ɑ- and β-actin (figure 4C-E). Free, not biotinylated sephin1 reduced actin retrieval (supplementary figure 3D-E). Given the interaction between sephin1 and β-actin, we investigated the impact of sephin1 on β-actin biochemical properties. First, we incubated purified human β-actin (250 nM) with sephin1 at different concentrations and analyzed actin partition via a sedimentation assay. By western-blotting, we found that sephin1 induced the precipitation of β-actin in a dose-dependent manner with an estimated EC50 of 150 μM (figure 6A-B). Sephin 1 precipitated efficiently α-actin (EC50= 215 μM) and had a weak effect on BSA (EC50= 790 μM, supplementary figure 4A-B). Differential centrifugation analysis indicated that sephin1 sedimentates β-actin already at 16k g (supplementary figure 4C-D). Altogether, our data supported the interaction between sephin1 and ɑ- and β-actin.

**Figure 3.**
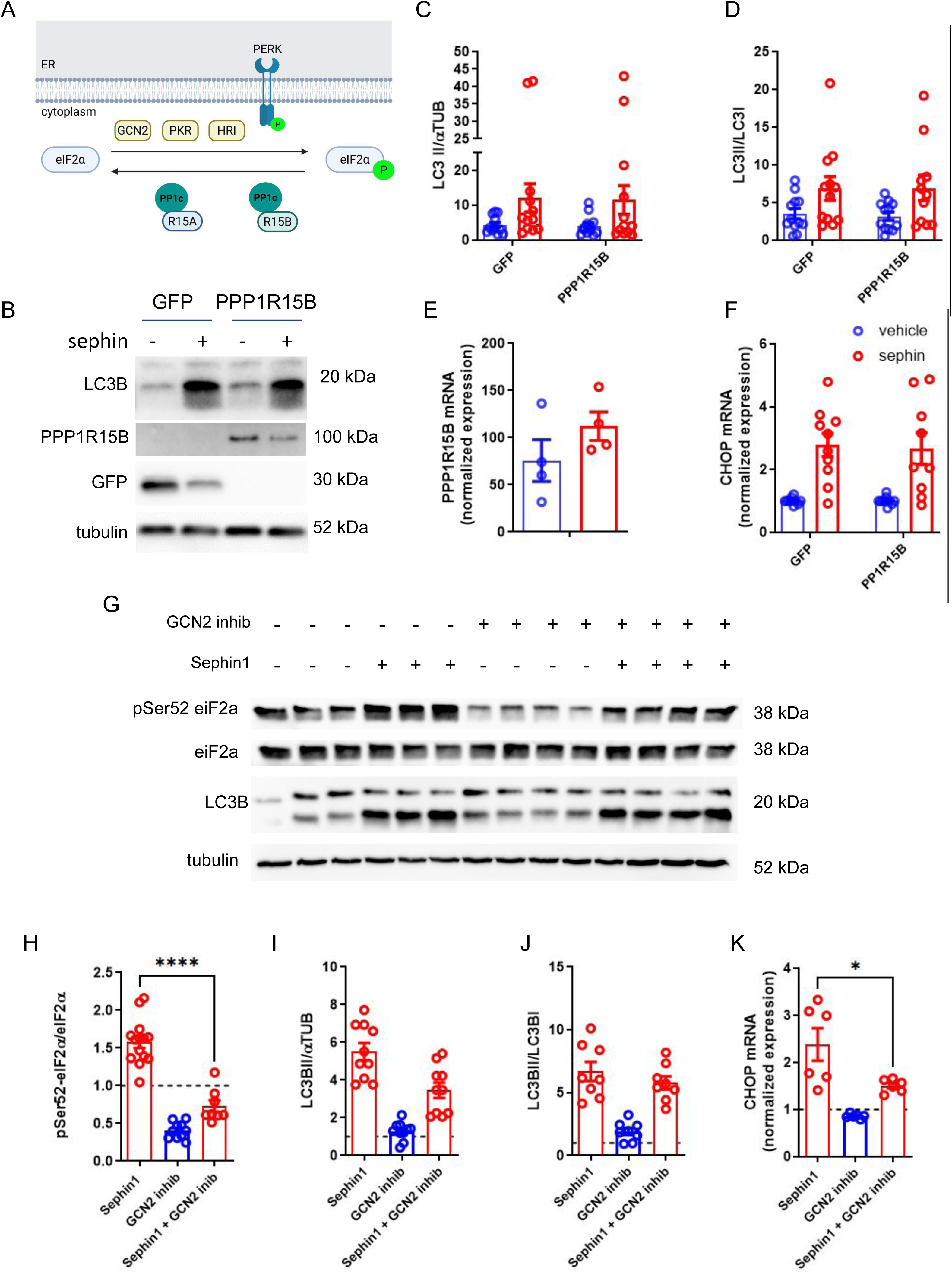
Sephin1 jeopardizes the autophagy flux via GCN2 kinase. (A) The phosphorylation of eIF2α is a key cellular response to stress that inhibits global protein synthesis. Various stresses, such as ER stress or nutrient deficiency, activate eIF2α kinases, such as PERK, PKR, HRI, and GCN2. The accessory protein PPP1R15A or PPP1R15B allow the protein phosphatase 1 (PP1) to dephosphorylate eIF2α and restore protein synthesis. (B) We monitored the impact on the autophagic flux in HeLa cells overexpressing GFP or PPP1R15B. (C-D) The graphs report the quantification of LC3B II levels, normalized to α-tubulin (C) and LC3II/LC3I ratio (D). Data are expressed as mean ± SEM, n =8-10. (E-F) qRT-PCR analysis of the levels of PPP1R15B (E) and CHOP (F) mRNAs. Data show 2^-△△Ct^ and are expressed as mean ± SEM, n =8-9. (G) We analysed eiF2α phosphorylation and LC3B conversion in cells treated with sephin1 alone (100μM, 2 hours) or in combination with the GCN2 inhibitor A92 (100 nM, 2 hours). (H-J) The graphs report eIF2a phosphorylation (H), LC3B II levels, normalized to α-tubulin (I), LC3II/LC3I ratio (J) and CHOP mRNA levels. Data show fold over vehicle and are expressed as mean ± SEM, n =6-9.

**Figure 4.**
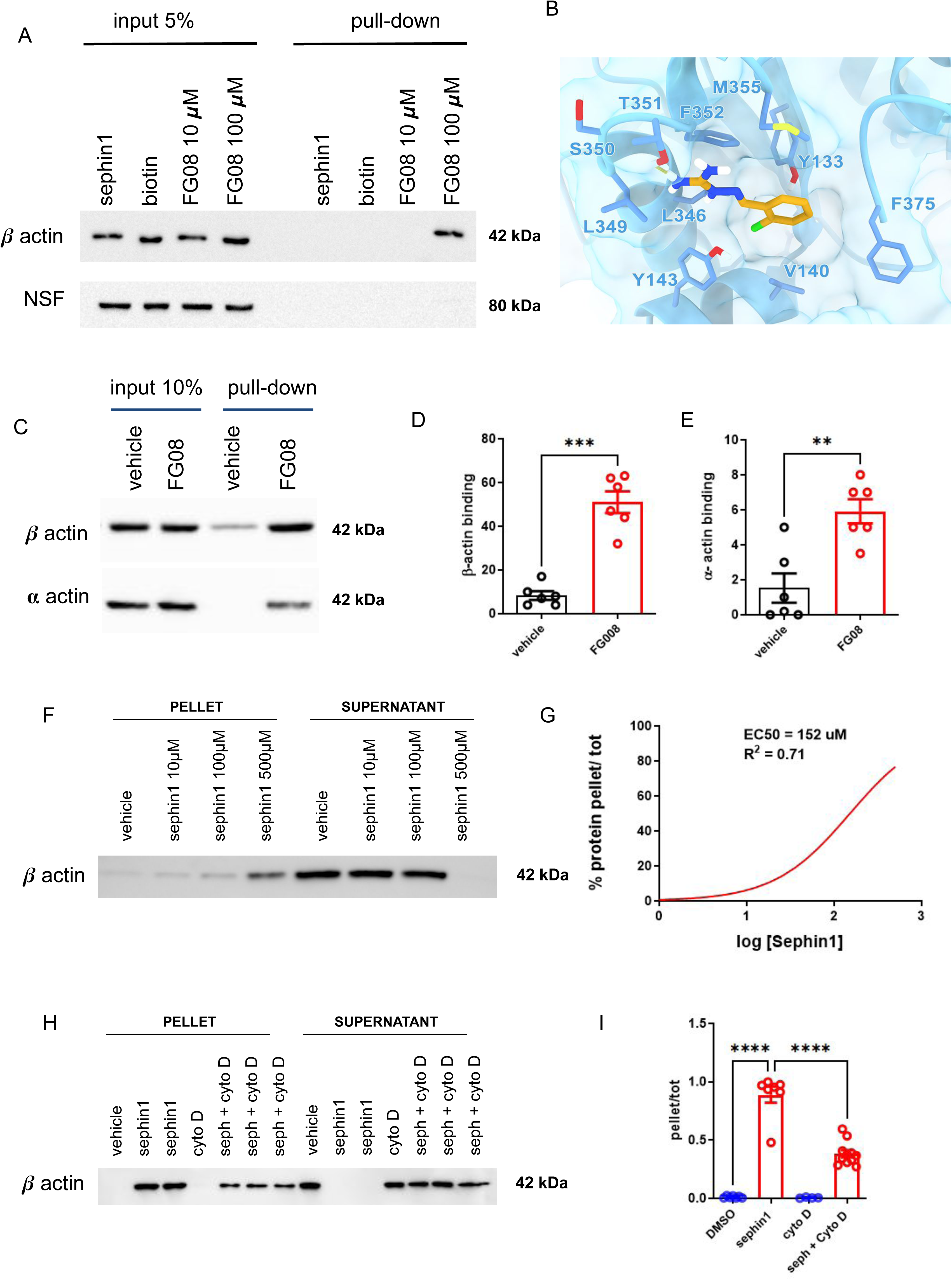
Sephin1 binds g-actin. (A) We used the biotinylated sephin1 derivative (FG08) to bind and isolate putative sephin1 targets. The pulldown assay suggested that sephin1 binds β-actin. An unrelated protein, NSF, was used as a control. (B) Molecular docking prediction of sephin1 binding site. (C) Sephin1 binds both purified α- and β-actin. (D-E) The graphs indicate the amount of bound protein, expressed as a fraction of the input. Data are expressed as mean ± SEM, n =6. (F) sephin1 induces β-actin precipitation upon centrifugation at 16k g. (G) The graph reports the non linear regression of the precipitation profile of purified β-actin. (H) We studied β-actin precipitation in presence of sephin1 alone or in combination with cytochalasin D (200 μM). (I) The graphs indicate the amount of precipitated protein, expressed as a fraction of the input. Data are expressed as mean ± SEM, n =7.

To identify the sephin1 binding site on β-actin, we combined photo-affinity labeling (PAL) chemistry with LC–MS/MS analysis. A photoactivatable sephin1 derivative (EC186; supplementary figure 5A) was synthesized and used to induce covalent crosslinking to actin upon UV irradiation. UV activation triggered nitrogen extrusion from the diazirine moiety, generating a highly reactive carbene intermediate capable of forming covalent bonds with proximal amino-acid side chains. High-resolution LC–MS/MS analysis of in-gel digested actin samples identified a modified peptide bearing a mass shift of +520 Da, consistent with the activated PAL-sephin1 adduct. The experimental precursor ion (m/z 521.14270) matched the theoretical exact mass (m/z 521.14226) with a mass error below 1 ppm, confirming the identity of the covalent adduct (supplementary figure 5B). MS/MS fragmentation localized the modification to the β-actin peptide QEYDESGPSIVHR (residues 360–372). To obtain atomistic insight into sephin1 binding, molecular docking calculations were performed. The search region was defined following an unbiased analysis of druggable sites in the cryo-EM structure of human G-actin (PDB: 8COG) using the SiteMap algorithm, with subsequent focus on regions proximal to the peptide identified by MS/MS (supplementary figure 5C). The most frequently predicted binding pose positioned sephin1 within a predominantly lipophilic pocket, where the chlorophenyl group establishes van der Waals interactions with residues L346, Y143, Y133, V140, and F375. In addition, the guanidyl-hydrazone group forms a hydrogen bond with the side chain of T351 (figure 4B and supplementary figure 5D). Notably, this predicted pocket overlaps with binding sites reported for other actin ligands, including cytochalasin D (supplementary figure 5E), suggesting potential competition between cytochalasin D and sephin1 for β-actin binding ^51–53^. Consistent with this hypothesis, cytochalasin D significantly reduced β-actin precipitation induced by sephin1 (figure 4H–I) and partially prevented Golgi disintegration observed following sephin1 treatment (supplementary figure 5F). Sephin1-induced actin sedimentation occurred rapidly (supplementary figure 6A), indicating that sephin1 interferes with actin dynamics, likely by reducing the pool of G-actin (supplementary figure 6C–D). Moreover, the polymerization rate of actin pre-incubated with sephin1 was significantly decreased (supplementary figure 6E–F). Given the confirmed interaction between sephin1 and β-actin, we further assessed the impact of sephin1 on actin biochemical properties. Actin sedimentation induced by sephin1 was completely abolished in the presence of 0.01% Triton X-100 (supplementary figure 7A–B). In conclusion, our data indicate that sephin1 binds directly to actin monomers, influences actin structural and biochemical properties, and its polymerization.

### Sephin1 alters the organization of actin cytoskeleton

Prompted by such biochemical evidence, we asked whether sephin1 modifies the actin cytoskeleton organization. We used Alexa 594-phalloidin to stain F-actin in HeLa cells treated with sephin1 (100 μM, 2 hours) or vehicles. We noticed that sephin1 promotes the formation of cytoplasmic actin clusters reminiscent of toxin induced actin patches/foci ^54^. These structures were transient in nature, since they almost disappeared after an overnight washout (supplementary figure 8A-B). The ARP2/3 complex is involved in shaping the actin cytoskeleton ^55^. We analyzed F-actin organization and LC-3 puncta frequency in the presence of sephin1 (100 μM, 2 hours) and of the ARP2/3 inhibitor CK-666 (250 μM, 2 hours). Sephin1 induced the appearance of actin clusters and LC-3 dots but the co-treatment with CK-666 significantly reduced the formation of them both (figure 5A-C and supplementary figure 8C). Next, we analyzed the autophagic flux in presence of sephin1 (100 μM, 2 hours) and CK-666 (250 μM, 2 hours). Strikingly, CK-666 reduced the impact of sephin1 on LC3-II accumulation (figure 5D-F).

**Figure 5.**
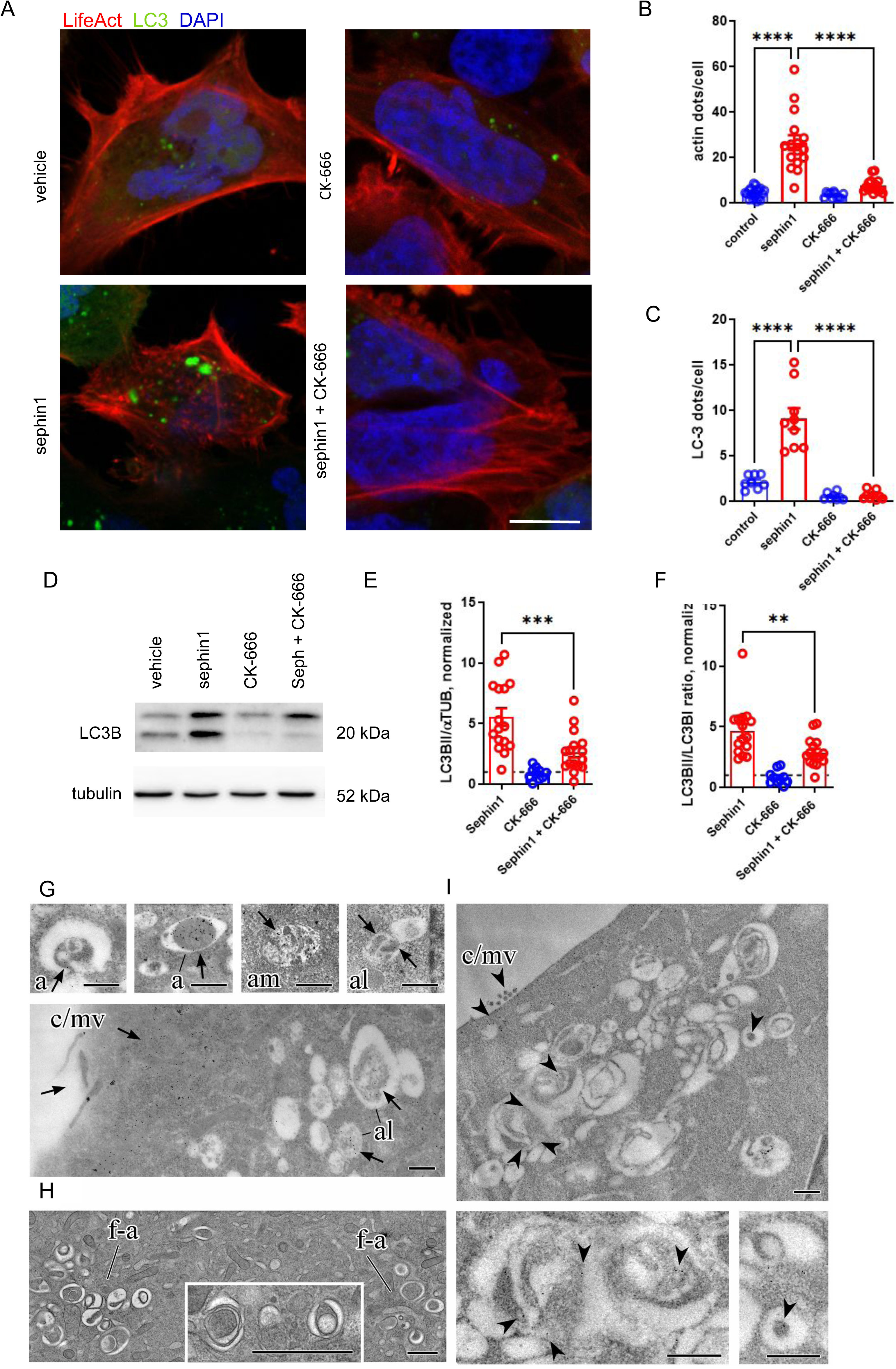
Sephin1 links the actin cytoskeleton to autophagy. (A) We monitored actin cytoskeleton and autophagic flux in HeLa cells expressing the lifeAct (red) and GFP-LC3 (green) reporters upon 2hours treatment with sephin1 (100 μM) or the ARP2/3 inhibitor CK-666 (250 μM) alone or in combination; scale bar= 5 μm. (B-C) The graphs report the number of actin (B) or LC3 positive dots (C). Data are expressed as mean ± SEM, n=8-10. (D) We assessed autophagy by monitoring LC3I and LC3B II levels in HeLa cells treated for 2 hours with sephin1 (100μM) in the presence or absence of CK-666 (250 μM). (E-F) The graphs report the quantification of LC3B II levels, normalized to α-tubulin (E) and LC3II/LC3I ratio (F). Data show fold over vehicle and are expressed as mean ± SEM, n =10-12. (G-I) Immunoelectron microscopic localization of actin and morphological features in cryofixed HeLa cells exposed for 2 hours either to sephin1 (100 μM) alone or together with CK-666 (250 μM). (G) Upon sephin1 treatment, distinct groups of several anti-actin immunogold particles (arrows) were regularly found onto the core of (forming) autophagosomes (a), but also on partially degraded contents of amphisomes (am) and autolysosomes (al) – in addition to robust labelling of cortical actin/microvilli (c/mv) and dispersed label throughout the cytoplasm; scale bar= 500nm. (H) Ultrastructure of various endomembrane compartments suggestive of abnormal, failed autophagosomes (f-a), characterizing Hela cells upon simultaneous administration of CK-666 (250 uM) and sephin1 (100 uM) for 2 hours; scale bars=1 µm. (I) Just sporadic, individual anti-actin immunogold particles (arrowheads) occur associated with putatively aberrant autophagosomes and Golgi derivatives in cells exposed for 2 hours to sephin1 (100 μM) together with CK-666 (250 μM) though moderate label is seen across cortical actin and microvilli, serving as internal control; scale bar= 500 nm.

EM together with immunogold labelling of endogenous actin and LC3 complemented these findings (figure 5G, supplementary figure 9). In cells treated with sephin1 (2 hours, 100 μM) we observed distinct labelling with several anti-actin immunogold particles on the cytoplasmic core of (forming) autophagosomes (figure 5G, supplementary figure 9C), resembling the actin enrichment inside autophagosomes shown in ^42^ and our controls. Some amphisomes and nascent autolysosomes showed internal actin label (figure 5G), whereas mature autolysosomes and terminal lysosomes were negative. Anti-LC3 clearly labelled autophagosomes and autolysosomes (supplementary figure 9B), as in starvation induced autophagy ^28^ and moderately also the swollen Golgi elements/derivatives (supplementary figure 9B). The simultaneous administration of CK-666 (250 μM) and sephin1 (100 μM) for 2 hours caused remarkable deformation of most autophagic organelles (Fig 5H), resembling failed autophagosomes ^42,56^. All aberrant organelles, including Golgi derivatives were rarely labeled by LC3, (supplementary figure 9B, arrowheads). Moderate actin label was observed, restricted to cortical actin and microvilli, but clearly not associated with autophagic organelles upon combined treatment with CK666 and sephin1 (figure 5I). Altogether, these results nominate actin as a bona fide target of sephin1 involved in the modulation of the autophagic flux.

### Actin clusterization precedes autophagy impairment and the ISR response

To gain an overview on the early events preceding LC3-II accumulation and ISR response, we investigated the cellular effect of sephin1 in a time-dependent manner. We studied actin organization, Golgi morphology, LC3-I and LC3-II accumulation, and eIF2 phosphorylation upon 5, 10, 30, 60, 120, and 240 minutes of sephin1 exposure through means of biochemistry and fluorescence microscopy (figure 6 and supplementary figure 10). Conspicuous actin clusters appeared already after 10 minutes of sephin1 treatment (figure 6A, C and supplementary figure 10A) and persisted during the next 120 minutes. The number of LC-3 positive fluorescent dots and corresponding protein levels also increased in this initial phase (figure 6B, C, supplementary figure 10C). LC3-II accumulated significantly further (figure 6D and supplementary figure 10E-H). Golgi integrity was severely compromised after 60 minutes (figure 6A, C, and 10B). CHOP mRNA levels rose after 120 minutes (supplementary figure 10D) and eIF2 phosphorylation increased overtly only after 240 minutes (supplementary figure 10E, F, and I). Notably, EM detected abundant phagophores and nascent autophagosomes, and first alterations of Golgi morphology already after 30 minutes of sephin 1 administration (supplementary figure 10J). Together, our data suggest that changing actin organization is the first event in a cascade that includes autophagic impairment, Golgi disintegration, and ISR activation.

**Figure 6.**
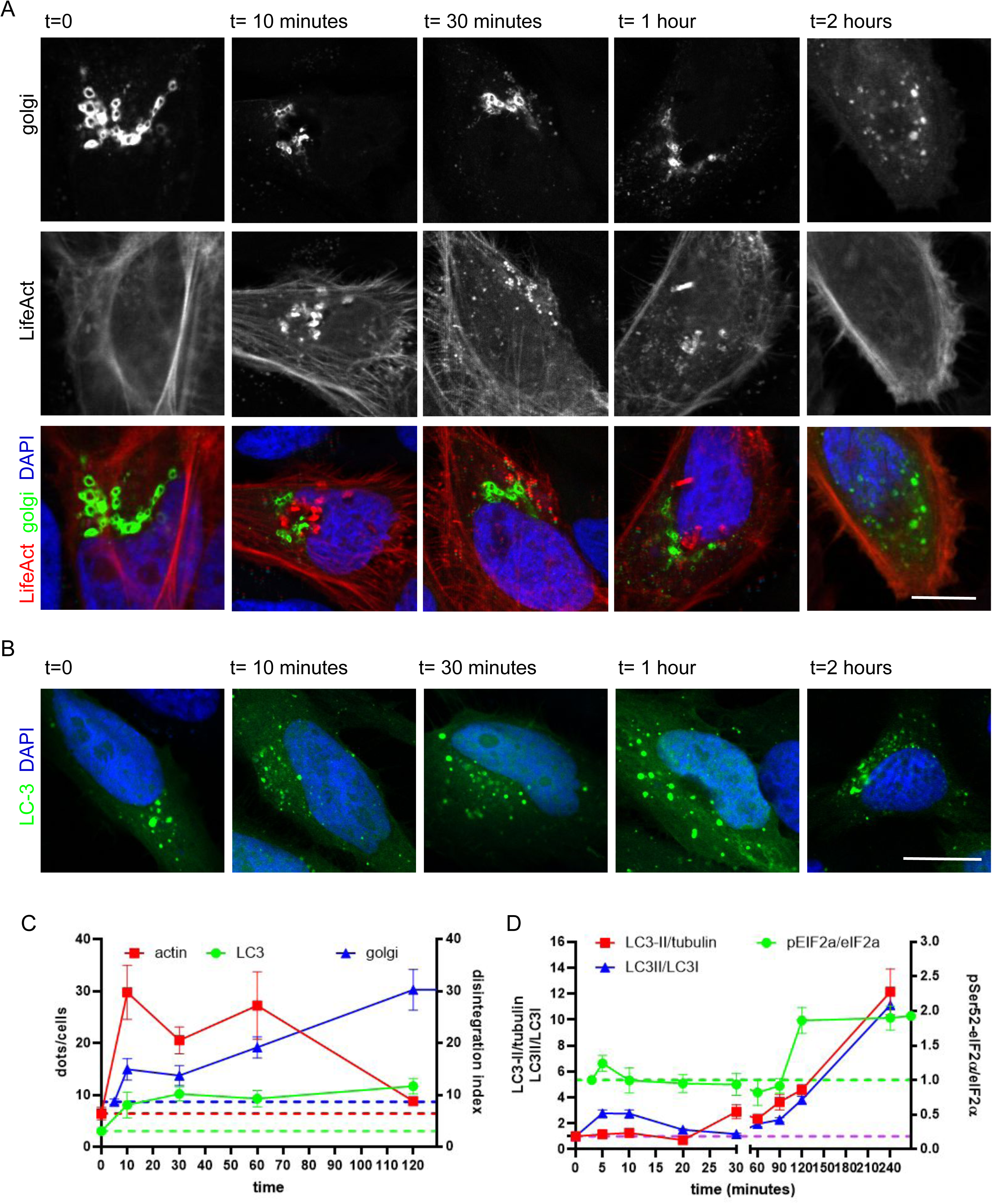
Analysis of sephin1 activity along time. (A) We studied Golgi morphology (GFP-Golgi), and actin cytoskeleton (LifeAct) in HeLa cells upon 10, 30, 60, or 120 minutes treatment with sephin1 (100 μM); scale bar= 5 μm. (B) We studied autophagy (GFP-LC3) in HeLa cells upon 10, 30, 60, or 120 minutes treatment with sephin1 (100 μM); scale bar= 5 μm. (C) The graph indicates the number of actin and LC3 positive dots as well as the Golgi disintegration index. Data are expressed as mean ± SEM, n=8-10. (D) The graph reports the quantification of LC3B II levels, normalized to α-tubulin, LC3II/LC3I ratio, and eIF2a phosphorylation. Data are expressed as mean ± SEM, n=5-6

### ALP stimulation is a long term consequence of sephin1 acute treatment

The mTOR-TFEB pathway is pivotal in regulating autolysosomal response ^57^. Under nutrient-rich conditions, mTORC1 is tethered to the lysosomal membrane where it promotes the cytoplasmic retention of TFEB via phosphorylation. Upon starvation, inactive mTORC1 leaves the lysosome and TFEB translocates to the nucleus ^58^. Consequently, we monitored by confocal imaging mTOR subcellular distribution in control conditions, upon sephin1 treatment, and in starvation. While in control condition the mTOR signal was concentrated in the perinuclear area, it became dispersed throughout the cytoplasm upon sephin1 treatment or starvation (figure 7A-B). Lysosomal mTOR phosphorylates several substrates, as 4EBP and P70s6K ^14^. Sephin1 treatment significantly impaired 4EBP and P70s6K phosphorylation (figure 7C-D). Finally, sephin1 and the mTOR inhibitor torin1 (1 μM, 2 hours) both promoted the nuclear translocation of TFEB (figure 7E-F). Sephin1-induced TFEB nuclear translocation was not consequent to oxidative stress or calcium release (supplementary figure 11A-B), nor did it require Mucolipin1 or calcineurin (supplementary figure 11C-E). Altogether, these observations suggest that sephin1 triggers TFEB nuclear translocation via a non-canonical pathway.

**Figure 7.**
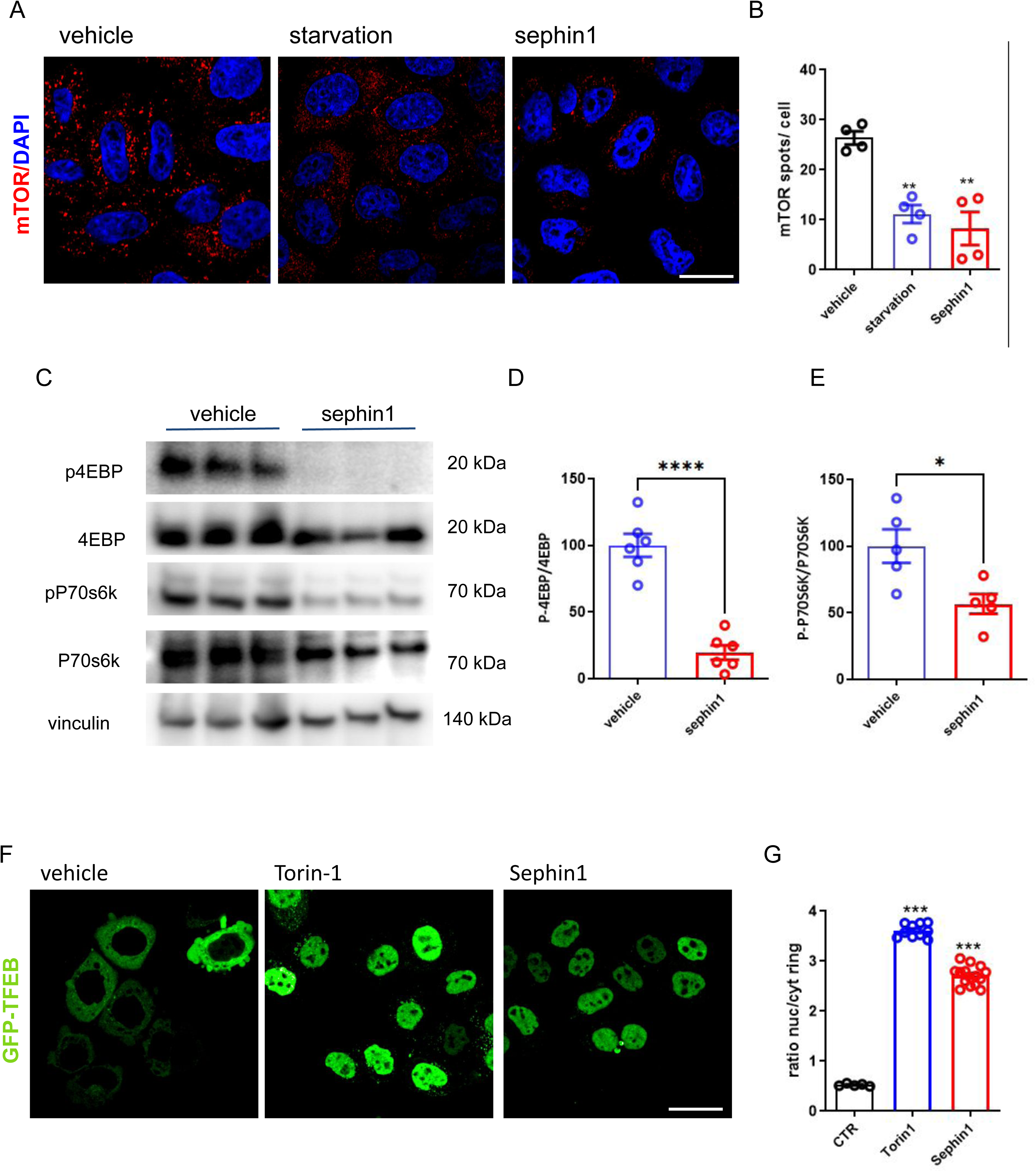
Sephin1 engages the mTOR-TFEB axis. (A) We monitored mTOR intracellular distribution in HeLa cells upon starvation or sephin1 treatment (100 μM); scale bar =10 μm. (B) The graph reports the number of mTOR spots. Data are expressed as mean ± SEM, n=4. (C) Biochemical analyses of the phosphorylation of the mTOR substrates 4EBP and P70s6K in HeLa cells treated with sephin1 (100 μM). (D-E) The graphs report 4EBP (D) and P70s6K (E) phosphorylation. Data are expressed as mean ± SEM, n=5. (F) We studied GFP-TFEB nuclear localization in HeLa cells upon 2 hours treatment with torin-1 (500 nM) or sephin1 (100 μM); scale bar=10 μm. (G) The graph reports GFP-TFEB nuclear localization, measured as nuclear to cytoplasm fluorescent signal ratio. Data are expressed as mean ± SEM, n=9.

By monitoring TFEB localization in a time-dependent manner, we noticed that sephin1-induced TFEB nuclear translocation peaked after 4 hours of wash-out (figure 8A-B). In the nucleus, TFEB stimulates the transcription of key ALP genes ^59^. TFEB-induced expression peaks 6-8 hours upon its nuclear translocation Accordingly, we monitored the levels of ATP6VOE1, ATP6V1h, P62, and RRAGD mRNA, four well established TFEB targets, right after acute sephin1 treatment (100 μM, 2 hours) or upon 2, 3, and 6 hours long wash out. Torin 1 (1 μM) served as positive control. Acute sephin1 treatment did not promote the expression of the four TFEB targets (figure 8C). Instead, we observed a partial effect upon 2- and 3-hours wash-out (supplementary figures 12A-B), and overt induction upon the 6 hours wash out (figure 8D). It is well known that, eventually, TFEB transcriptional activity increases the number and the activity of lysosomes ^59^. By the DQ-BSA assay, we assessed the lysosomal activity in cells treated with sephin1 or torin1 following a 6-hours wash-out after 2 hours long treatment. Indeed, sephin1 correlated with an increase of DQ-BSA fluorescent spots, witnessing an increase in lysosomal activity after the release of the sephin1-induced autophagy blockade (figure 8E-F). Since translation inhibitors also activate TFEB ^60^, we treated HeLa cells with cycloheximide (100 μg/mL, 2 hours) but we did not observe any major impact on the transcription of TFEB targets after the 6 hours long wash out (supplementary figure 12D). Altogether, our data suggest that sephin1 pulse stimulates lysosomal activities via the mTOR-TFEB pathway. TFEB links lysosomal biogenesis and autophagy ^15^. As such, we monitored the autophagy flux in HeLa cells treated with sephin1 (100 μM, 2 hours) followed by 8 hours wash out and exposed for further 2 hours to DMSO or bafilomycin (100 nM). The quantification of LC3-II protein levels showed that the initial sephin1 pulse significantly stimulated autophagy (figure 9A-B), in absence of overt ISR (supplementary figure 12C). Finally, the well-established autophagic inducer trehalose did not elicit any major response in the same condition (supplementary figure 12C-E). Next, we wondered whether the remodeling of the actin cytoskeleton taking place in the acute phase may affect the late-stage autophagic induction. At this aim, we treated HeLa cells with sephin1 (100 μM, 2 hours) alone or in combination with CK-666 (250 μM, 2 hours). After the wash-out and the treatment with DMSO or bafilomycin (100 nM, 2 hours), we measured by western-blotting LC3-II. Noteworthy, the co-treatment with CK-666 in the acute phase significantly blunted sephin1 activity (figure 9C-D). EM analyses as well provided strong morphological evidence for efficient autophagy following a sephin1 pulse and subsequent cell recovery in normal medium (figure 9E). Upon sephin1 administration (100 μM, 2 hours) we found conspicuous enrichment of autophagic organelles and Golgi alterations, those almost disappeared during the 6 hours long wash-out. BafA administration to cells subjected to a 2 hours sephin1 pulse, followed by overnight wash out, revealed accumulation of abundant autophagic organelles. By contrast, cells exposed to ARP2/3 inhibitor CK-666 (250 μM) together with sephin1 during the pulse phase remained almost free of autophagic organelles upon wash out and BafA administration. Conclusively, we propose that the initial actin cytoskeleton remodelling induced by sephin1 is instrumental for the late autophagic response.

**Figure 8.**
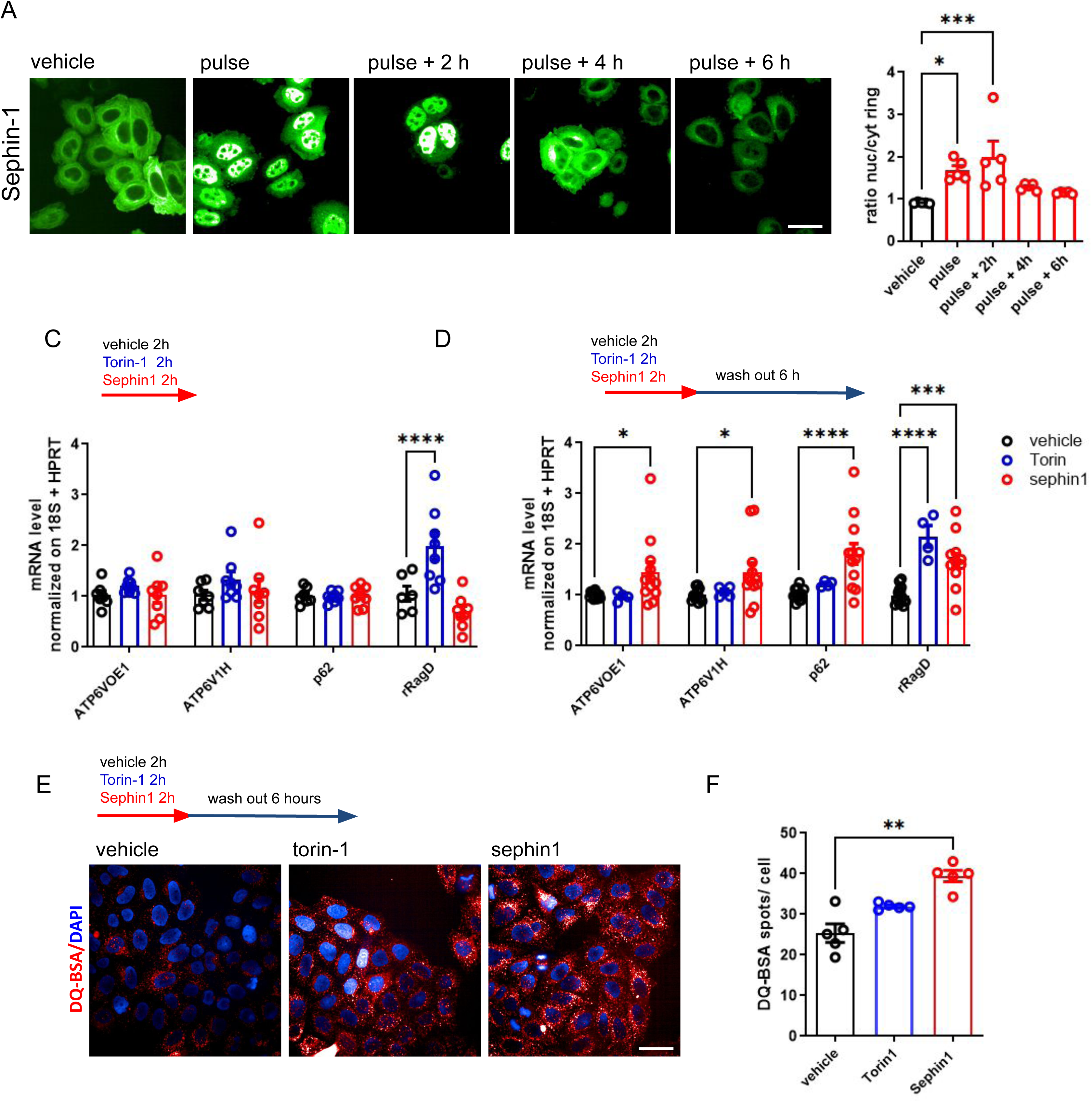
Sephin1 pulse induces a late autophagic response. (A) We studied GFP-TFEB nuclear localization in HeLa cells following an initial 2 hours treatment; scale bar =10 μm. (B) The graph reports GFP-TFEB nuclear localization, measured as nuclear to cytoplasm fluorescent signal ratio. Data are expressed as mean ± SEM, n=50 cells, from 3 independent experiments. (C-D) We treated HeLa cells for 2 hours with torin-1 (500 nM) or sephin1 (100 μM). We processed the cells right after the treatment or upon a 6 hours long wash-out. qRT-PCR analysis of the levels of key TFEB targets right after 2 hours treatment with sephin1 (C, 100 μM) or upon 6 hours long wash-out. Data show 2^-△△Ct^ and are expressed as mean ± SEM, n =6-10. (E) We assessed lysosomial activity by measuring DQ-BSA fluorescence in HeLa cells after the pulse-wash out protocols with torin-1 or sephin1; scale bar =10 μm. (F) The graph reports the number of DQ-BSA spots. Data are expressed as mean ± SEM, n=5.

**Figure 9.**
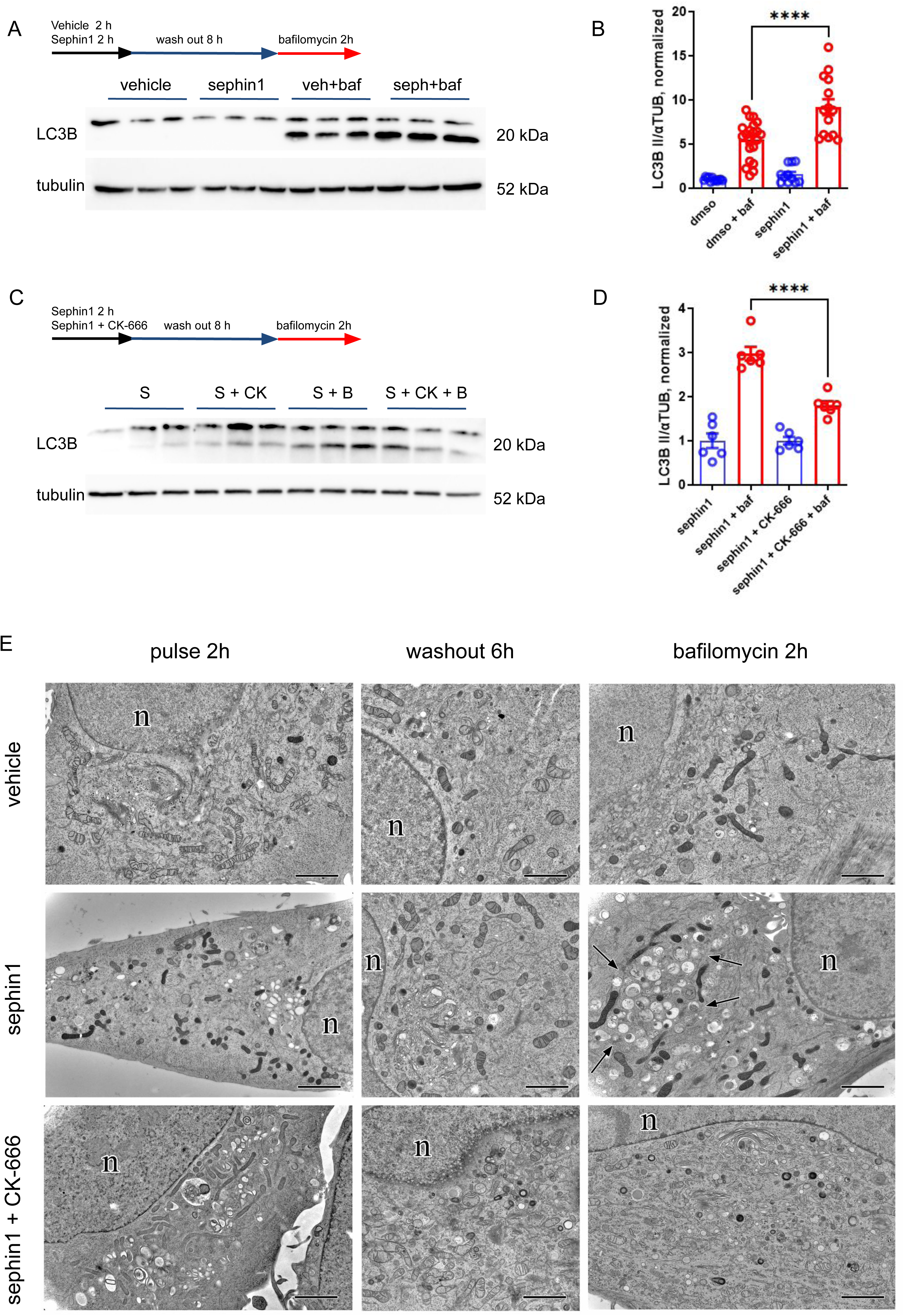
Sephin1 pulse induces a late autophagic response via actin. (A) We assessed autophagy by monitoring LC3B II levels in HeLa cells treated for 2 hours with sephin1 (100 μM) or vehicle, followed by 8 hours long wash out and eventually exposed to bafilomycin 1 (baf, 100 nM 2 hours). (B) The graph reports the quantification of LC3B II levels, normalized to α-tubulin; data are expressed as mean ± SEM, n =10-12. (C) We monitored LC3B II levels in HeLa cells treated for 2 hours with sephin1 (100 μM, 2 hours) alone or in combination with CK-666 (250 μM, 2 hours), followed by 8 hours long wash out and eventually exposed to bafilomycin 1 (baf, 100nM 2 hours). (D) The graph reports the quantification of LC3B II levels in bafilomycin 1 condition, normalized to α-tubulin, and expressed as fold over not bafilomycin 1 condition; data are expressed as mean ± SEM, n =6. (C) EM micrographs of cryofixed HeLa illustrating long term effects upon cell exposure to vehicle, sephin1 alone (100 μM, 2 hours), or in combination with actin inhibitor CK666 (250 μM, 2 hours). A 6 hours long drug wash out rescues any ultrastructural abnormalities and organelle jam observed upon incubation with sephin1 or sephin1 plus CK-666. However, only in the cultures previously exposed to the sephin1 pulse, bafilomycin 1 administration (100 nM, 2 hours) reveals a strong late autophagic response, as indicated by abundant autophagic organelles (arrows). Co-administration of CK666 during the pulse phase prevents the late autophagic response. (g)= Golgi/TGN, (n)=nucleus; scale bars= 2 μm.

**Figure 10.**
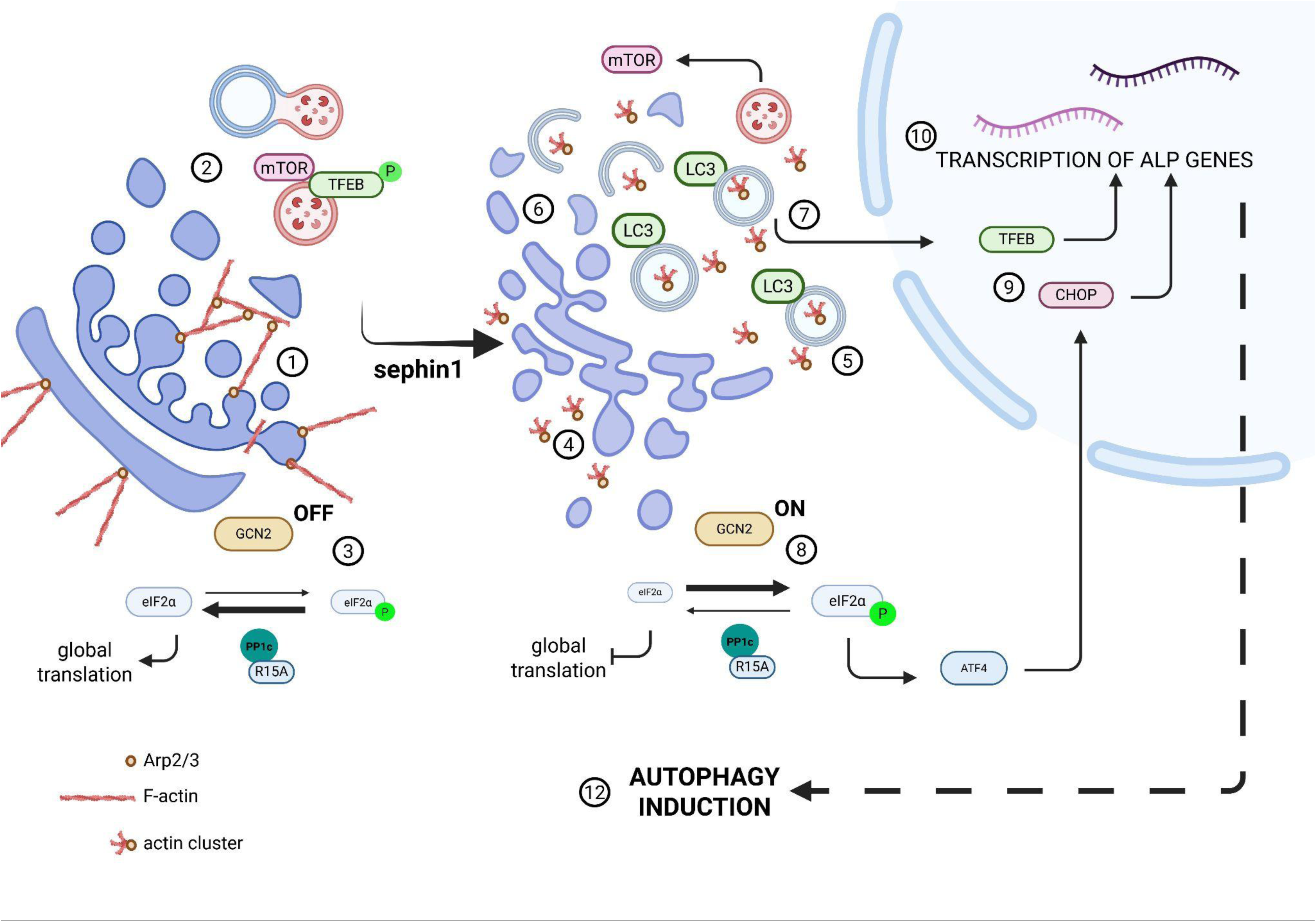
Sephin1 pulse induces a late autophagic response via actin. We propose that sephin1 induces autophagy as a late response to an initial actin cytoskeleton remodelling. (1) Actin cytoskeleton contributes to maintain Golgi integrity, allows Golgi scission, and drives vesicle budding. (2) The autophagy flux is functional and mTOR tethers TFEB on the autolysosomal membrane. (3) GCN2 is inactive and eIF2 supports global translation. (4) Sephin1 induces the formation of cytoplasmic F-actin clusters via Arp2/3. (5) Actin clusters initiate the formation of autophagomes. (6) The alteration of cytoplasmic actin cytoskeleton causes Golgi damage. Golgi disintegration procures vesicles contributing to the assembly of a large pool of immature autophagosomes. (7) Autophagic impairment inhibits mTOR, stimulates its cytoplasmic relocalization thus promoting the nuclear translocation of TFEB. (8) Actin remodelling triggers GCN2 activation and, eventually, eIF2 phosphorylation. (9) eIF2 phosphorylation promotes the non-canonical translation of ATF4 and the consequent CHOP induction. (10) TFEB and CHOP stimulate the transcription of ALP genes. (11) The activation of both eIF2-CHOP and mTOR-TFEB pathways allow the activation of a functional autophagic response.

## Discussion

The mechanism of action of sephin1 is highly controversial. Originally, sephin1 and the structurally related GBZ were described as selective inhibitors of the PPP1R15A subunit. As such, sephin1 should impair the PPP1R15A-mediated P-eIF2α dephosphorylation ^17,18,61^. Further studies challenged the previous observations and argued that PPP1R15A is neither binding GBZ nor sephin1 ^24,25^. Hence, sephin1 target and mechanism of action are far from being fully assessed. Our data indicate that sephin1 binds β-actin in a pocket overlapping with the cytochalasin-D binding site and influences its structural features. Our cell-free studies suggest a weak interaction with G-actin, as suggested by the EC50 around 150 μM in the precipitation assay and the binding happening above 100 μM sephin1 concentration. In vitro works on polymerization kinetics posit the critical concentration of G-actin in the high nanomolar range ^62^. We noticed that the incubation with sephin1, either previous or contextual, severely reduced the polymerization rate of G-actin. As such, it is well possible that our biochemical assays underestimated the affinity between sephin1 and G- or F-actin. Accordingly, we did monitor cellular effect at lower concentration, starting from 50 μM. Cytochalasin-D binds G- and F-actin with largely different affinity (Kd F-actin: 2 nM; Kd G-actin 20 μM), suggesting that the quaternary structure of actin affects its binding with small molecules ^63^.Thus, we may not exclude that in the cellular context sephin1 binds actin with higher affinity.

Our data suggest that sephin1 induces the assembly of large actin structure, such structures encompass at least three G-actin monomers, as they are recognised by phalloidin, and interact with Arp2/3. The Arp2/3 complex enhances actin filament nucleation, cross-linking, and branching ^64^. Interestingly, Mi et al., already reported that branched actin structures shaped by Arp2/3, cofilin, and CapZ are present inside the cavity of forming autophagosomes (phagophores/isolation membranes). The presence of branched actin network inside the phagophore is required for autophagosome formation, as the treatment with the Arp2/3 inhibitor CK666 prevents the proper curvature of the phagophore and causes the collapse of omegasomes and phagophores, eventually blocking autophagy ^42^. It is tempting to speculate that sephin1 promotes the formation of Arp2/3 dependent structures similar to the one described by Mi and colleagues. Indeed, we identified endogenous actin by immuno-EM inside structurally normal, forming autophagosomes in sephin1 treated cells but not in any kind of (deformed) autophagic organelles occurring upon co-treatment with sephin1 and CK-666. Sephin1-driven actin structures may be sufficient to promote the initial formation of autophagosomes but not yet to trigger further autophagy progression and waste clearance in an early phase. The abundant (forming) autophagosomes we observed upon acute sephin1 treatment might not be fully decorated by the V-ATPase, as suggested by the lack of responsiveness to Bafilomycin-1 revealed by biochemistry. They, could, therefore, represent immature autophagosomes. Our molecular analyses clearly indicate that V-ATPase subunits mRNAs transcription increases not immediately upon sephin1 pulse, but after 6 hours. One possible explanation is that sephin1 triggers the formation of branched actin structure resembling the ones described by Mi and colleagues, supported by Arp2/3, but missing the complete protein network instrumental to support autophagy progression in the short term.

Actin cytoskeleton is critical for Golgi dynamics, mechanics, and morphology ^65,66^. In particular, Golgi-actin interaction requires Arp2/3-driven actin projections ^67^. By interacting with WHAMM, Arp2/3 becomes essential in maintaining the Golgi integrity and facilitating membrane transport ^68^. We clearly observed a transient Golgi disintegration upon sephin1. Sephin1-induced actin clusters in the cytoplasm may impact on the local actin cytoskeleton, putatively influencing F-actin stability by reducing G-actin pool and/or sequestering Arp2/3 complexes, and eventually, harming Golgi integrity. The Golgi complex is a central hub within the endomembrane trafficking system. Although the ER is implicated as the primary membrane source necessary for the formation of autophagosome precursors in canonical autophagy ^69^, the Golgi is involved in the subsequent autophagosome elongation, and Golgi-mediated trafficking provides membrane components for autophagosome biogenesis. The Golgi also makes available pivotal autophagy regulators, including the protein autophagy-related (ATG)-9A ^46^. Strikingly, Golgi fragmentation can increase autophagosome biogenesis by feeding ATG-9A positive fragmented Golgi membranes ^70^. Similarly, Golgi fragmentation induced by prolonged exposure to low doses of Brefeldin A favours autophagosome biogenesis and induces the accumulation of autophagosomes ^71^. Indeed, Golgi membrane-associated degradation (GOMED) pathways generate autophagosomes morphologically like the canonical ones ^72,73^. As such, the accumulation of a heterogenous population of immature, possibly aberrant (forming) autophagosomes that we observed upon sephin1 acute treatment may well follow the transient disruption of the Golgi complex. However, our findings show that the sephin1 related Golgi fragmentation itself is not sufficient to stimulate autophagy in the long term. We did observe Golgi fragmentation and definitely aberrant autophagosomes in cells treated with sephin1 and CK-666. In those conditions we did report the strong late phase autophagic induction that follows the sephin1 only pulse. As such, Golgi fragmentation may support the sephin1 mechanism but does not explain it by itself.

In agreement with the original findings linking sephin1 to eiF2 phosphorylation, we reported that sephin1 indirectly increases eIF2α phosphorylation in a ER-stress independent manner, as we did not observe overt alterations in the levels of BiP and spliced-Xbp1 mRNAs or PERK activation upon sephin1 treatment. Among the kinases acting on eIF2α to reduce the proteostatic stress, GCN2 answers not only to a reduction of the amino acids outflows from the lysosome ^74^, but also to actin cytoskeleton remodelling. GCN2 monitors the state of the actin cytoskeleton and modification of the actin cytoskeleton elicits a GCN2-mediated eIF2α phosphorylation response ^75^. In turn, eIF2α phosphorylation increases the expression of the transcription factor CHOP via ATF4. We noticed that GCN2 inhibition was sufficient to abolish sephin1 induced eIF2 phosphorylation and CHOP induction but did not prevent autophagosome accumulation. As such, we hypothesize that sephin1 triggered F-actin clusterization and, besides its impact on Golgi, activated the GCN2-eIF2α-CHOP pathway. Strong evidence links the eIFα/ATF4 pathway to autophagy ^76,77^. Indeed, the GCN2 and PERK kinases and the transcription factors ATF4 and CHOP are required to increase the transcription of genes implicated in the formation, elongation and function of the autophagosome, such as p62, Atg7, Atg10 ^7^. However, the link between autophagy and CHOP is not straightforward, as CHOP limits autophagy in case of prolonged stress conditions ^78,79^.

mTORC1 is active on the lysosomal surface and responds to different stimuli, including growth factors, nutrients, energy status, and stressors ^80–82^. mTORC1 phosphorylates TFEB on serine 211, thus providing a binding site for the cytoplasmic 14-3-3 chaperones. The 14-3-3 proteins sequester TFEB in the cytoplasm ^83–85^. The inhibition of mTORC1 results in ALP induction upon TFEB dephosphorylation and its nuclear translocation. Our data clearly indicate that sephin1 promotes TFEB nuclear translocation. TFEB phosphorylation is tightly controlled by multiple mechanisms, including lysosomal calcium current and redox state. Lysosomal Ca2+ flux through mucolipin 1 (MCOLN1) stimulates calcineurin, which dephosphorylates TFEB. ROS activate TFEB indirectly via MCOLN-1-mediated lysosomal Ca2+ release and directly via oxidation of its key cysteine 212 ^86^. However, we excluded that sephin1 promotes TFEB translocation via either MCOLN1-calcineurin or ROS.

The accumulation of immature autophagosomes has a complex and dynamic influence on the mTORC1-TFEB pathway. Interestingly, not only the failed clearance of waste-loaden autolysosomes is harmful; also, the excess accumulation of autophagosomes subsequently unfused to lysosomes is cytotoxic, causing an energy deficit and elevates ROS production ^87^. Eventually, an autophagy flux blockade impairs the aminoacid content in the lysosomes. The Rag GTPase Ragulator complex couples amino acid availability to lysosomal recruitment and activation of mTORC1.When amino acids are abundant, Rag stimulates the recruitment of mTORC1 to the lysosomes. When the amino acid content turns low, Rag GTPases are inactive and so is, consequently, also mTORC1. Acutely, sephin1 jeopardizes the autophagic flux. Genuine (terminal) lysosomes became rare during sephin1-induced autophagy blockade. As such, sephin1 may reduce the amino acid flux through the lysosomes and thus promote TFEB translocation via Rag GTPases. Similarly, an altered autophagosome membrane composition may influence TFEB activation by changing the balance of factors that control its phosphorylation and translocation ^88^.

Notwithstanding the yet obscure mechanisms involved, our data clearly show that sephin1 promotes TFEB translocation. Indeed, the activation of TFEB is central in the signaling cascade that we proposed but is not sufficient to induce autophagy *per se*. Dang et al reported that three different translation inhibitors (i.e., CHX, Lactimidomycin, and Rocaglamide A) also elicit the nuclear translocation of TFEB, increasing autophagosome biogenesis, but do not support a functional autophagy ^60^. Thus, we speculate that the contribution of P-eIF2α-CHOP signaling is of paramount importance to promote a functional autophagic process. Our hypothesis is that sephin1 modifies a specific subset of the cytoplasmic actin pool, resulting in the accumulation of immature autophagosomes, associated with transient Golgi disintegration. Actin remodelling and acute autophagy flux blockage activates two distinct pathways, GCN2-eiF2-ATF4 and mTORC1-TFEB, which ultimately result in an increased autophagy.

Overall, our data contributes to the accumulating evidence that actin regulates multiple steps in autophagy. Interestingly, Chen and collaborators indicated G-actin as an important cofactor that stabilizes the PPP1R15-PP1 complex. The PPP1R15-PP1-G-actin ternary complex accelerates P-eIF2α dephosphorylation ^89^. As such, sephin1 may affect PPP1R15-PP1 activity indirectly via G-actin. Indeed, drugs impacting the actin cytoskeleton can affect eIF2α phosphorylation. Jasplakinolide binds G-actin and it has been proposed to increase eIF2α phosphorylation via lowering the free G-actin content ^90^. Similarly, sephin1 may reduce the cytoplasmic pool of available G-actin thus jeopardizing the formation of the PPP1R15-PP1-G-actin ternary complex. Overall, sephin1 may slow eIF2α dephosphorylation by binding to G-actin and preventing the formation of the active ternary complex. Consequently, our study does not exclude a possible, non-autophagy related, alternative effect of sephin1 on PPP1R15A and reinforces the role of actin cytoskeleton as a factor that regulates ISR.

Conclusively, our findings support a model where a transient, Sephin1-induced cytoskeletal and organelle stress event acts as a pharmacological stimulus. This acute phase, characterized by GCN2 activation and an initial autophagic block, is required to induce the subsequent nuclear translocation of TFEB and the enhancement of lysosomal biogenesis and autophagy. This biphasic MoA explains why Sephin1 is therapeutically beneficial in long-term disease models despite its apparent acute impact on organelle integrity.

