## supplementary figures for "Sephin1 rewires proteostasis through actin-dependent signaling"

**Supplementary figure 1.** (A) We assessed the viability of HeLa cells upon 2 hours treatment with sephin1 at different concentrations by the means of MTT assay. Data are expressed as mean ± SEM, n =10-12. (B) We assessed autophagy by monitoring LC3I and LC3B II levels in HeLa cells treated for 2 hours with sephin1 at different concentrations in the presence or absence of Bafilomycin 1 (BafA1, 100nM). (C-D) The graphs report the quantification of LC3B II levels, normalized to α-tubulin (B) and LC3II/LC3I ratio. Data are expressed as mean ± SEM, n =6-10. (E-F) Hemilogarithmic dose-response curve plotting LC3B II levels, normalized to α-tubulin (E) and LC3II/LC3I ratio (F). (G) We monitored lysosomal damage by staining with anti galectin-3 antibody. HeLa cells were cultured in fed condition or treated with LLOME (1 mM , 2 hours) or sephin1 (100µM, 2 hours), scale bar 20 µm. (H) The graph reports the number of galectin + spot per cell. Data are expressed as mean ± SEM, n =6.

**Supplementary figure 2.** (A) We monitored protein translation by means of puromycin labelling assays (10 µg/mL) in HeLa cells treated with vehicle (DMSO), sephin1 (100µM), or cycloheximide (CHX, 100 µg/mL). (B) The graphs report the quantification of the puromycin signal, normalized to ponceau. Data are expressed as mean ± SEM, n =5-6. (C) In the same experimental conditions, we assessed autophagy by monitoring LC3I and LC3B II levels. (D-E) The graphs report the quantification of LC3B II levels, normalized to α-tubulin (D) and LC3II/LC3I ratio (E). Data are expressed as mean ± SEM, n =5-6. (F) We assessed autophagy by monitoring LC3I and LC3B II levels in HeLa cells treated for 2 hours with sephin1 (100 µM) in the presence or absence of cycloheximide (CHX, 100 µg/mL). (G-H) The graphs report the quantification of LC3B II levels, normalized to α-tubulin (G) and LC3II/LC3I ratio (H). Data show fold over vehicle for sephin1 treatment or CHX for sephin1 + CHX treatment and are expressed as mean ± SEM, n =7-8.

**Supplementary figure 3.** (A) Structure of FG08, a biotinylated derivative of sephin1 (red box). (B) We incubated HeLa cell lysate with FG08 at 10 or 100 µM. Upon immobilization on streptavidin coated beads, we analysed putative interactors by SDS-PAGE followed by silver-staining. We analysed by MS/MS the band indicated by the arrow. (C) MS/MS spectra of the isolated band identified the protein as actin. (D) Sephin1 competes with FG08 for the binding to β-actin. We incubated purified β-actin with FG08 (100 µM) alone or in presence of 100 or 500 µM sephin1. (E) The graph indicates the amount of bound protein, expressed as a fraction of the input. Data are expressed as mean ± SEM, n =6.

**Supplementary figure 4.** (A) Sephin1 induces α-actin precipitation upon centrifugation at 16k g but does not influence albumin solubility. (B) The graph reports the non linear regression of the precipitation profile of purified α-actin and albumin. (C) We studied the impact of sephin1 on the precipitation profile of β-actin upon centrifugation at 16k or 100k g. (D) The graph indicates the amount of precipitated protein, expressed as a fraction of the input. Data are expressed as mean ± SEM, n=4.

**Supplementary figure 5.** (A) Structure of the PAL derivative of sephin1 EC186, and predicted structure and MW post UV irradiation. (B) MS/MS analysis of the sephin1 PAL derivative (upper graph, experimental peaks; lower graph, expected peaks). (C) MS/MS analysis of the actin peptide tagged by the PAL derivative. Uncertainty remains regarding the site of conjugation, either a tyrosine or an aspartic acid. (D) The model depicts the position of the target peptide identified by the MS/MS approach (red). Considering the structure of the PAL derivative, the binding pocket identified by in silico docking is compatible with the target peptide identified. (Inset) Atomic description of sephin1 binding pocket. (E) The putative binding pocket of sephin1 (yellow) overlaps with the site bound by cytochalasin D (green). (F) We monitored Golgi morphology by expressing the GFP-Golgi reporter in HeLa cells upon 2 hours of treatment with sephin1 (100 μM), cytochalasin-D (10 μM) either alone or in combination, scale bar= 5 μm.

**Supplementary figure 6.** (A) Time-course analysis of the precipitation profile of purified β-actin in presence of sephin1 (100 μM). (B) The graph indicates the amount of precipitated protein, expressed as a fraction of the input. Data are expressed as mean ± SEM, n=3. (C) Polymerisation rate of 2 μM pyrene-actin in presence of increasing amount of sephin1. (D) Linear regression of the initial 1000 seconds of the polymerisation curves. (D) Polymerisation rate of 2 μM pyrene-actin upon overnight incubation with increasing amount of sephin1. (E) Linear regression of the initial 1000 seconds of the polymerisation curves. Data are expressed as mean ± SEM, n=4.

**Supplementary figure 7.** (A) We studied the precipitation profile of purified β-actin in presence of sephin1 alone (100 μM) or in combination with 0.01 % Triton-X100. (B) The graph indicates the amount of precipitated protein, expressed as a fraction of the input. Data are expressed as mean ± SEM, n=6.

**Supplementary figure 8.** We monitored the actin cytoskeleton by means of phalloidin staining in HeLa cells treated for 2 hours with sephin1 (100 μM) or upon overnight washout (WO); scale bar= 10 μm. (B) The graph reports the number of actin-positive dots. Data are expressed as mean ± SEM, n=8-10. (C) Phalloidin staining in HeLa cells upon 2hours treatment with sephin1 (100 μM) or the ARP2/3 inhibitor CK-666 (250 μM) alone or in combination; scale bar= 10 μm.

**Supplementary figure 9.** Immunoelectron microscopic localization of endogenous actin and LC3, as well as ultrastructural features in HeLa cells exposed for 2 hours either to sephin1 (100 μM) alone or combined with CK-666 (250 μM), and processed through means of Tokuyasu's technique or cryofixation. (A) Sporadic, patchy accumulations (pc) of 6-7nm-wide, actin-positive filaments (arrows), potentially correlating with the fluorescent phalloidin-positive clusters. Pink lines mark corresponding areas, the dotted line the cluster's entire extension; scale bar=1μm. Distinct actin accumulations attached to the organelles' surface possibly corresponding to transient "autophagosomal actin comet tails", seen in live cell imaging experiments (DOI: 10.1016/j.cub.2015.05.042), were rarely caught in our EM-snapshots. (B) Anti-LC3 labelling of cryofixed Hela cells. Immunogold label (arrows) is seen on autophagosomes (a) and autolysosomes (al), and, moderately, also Golgi (derivatives: g) in cells treated with Sephin for 2 hours. Combined administration of CK-666 and sephin1 yielded just scarce, singular gold particles (arrow-heads) across endomembranes; scale bar=1μm. (C) Methodological controls for the localization of actin and LC3 (arrows) on forming autophagosomes (a); scale bar=1 μm.

**Supplementary figure 10.** (A-D) The graphs indicate the number of actin (A) and LC3 (B) positive dots as well as the Golgi disintegration index (C) and CHOP mRNA induction in HeLa cells upon 10, 30, 60, or 120 minutes treatment with sephin1 (100 μM). Data are expressed as mean ± SEM, n=6-10. (E-F) Biochemical analyses of eIF2 phosphorylation and LC3 conversion in HeLa cells treated with sephin1 (100 μM) up to 240 minutes. (G-I) The graph reports the quantification of LC3B II levels, normalized to α-tubulin (G), LC3II/LC3I ratio (H), and eIF2a phosphorylation (I). Data are expressed as mean ± SEM, n=5-6. (J) EM of cryofixed HeLa shows that administration of sephin1 (100 μM) for 30 minutes yields numerous (forming) autophagosomes (a) and locally alterations of the Golgi/TGN (g); scale bar =500 nm.

**Supplementary figure 11.** (A) We studied GFP-TFEB nuclear localization in HeLa cells upon 2 hours treatment with sephin1 alone (100 μM), in combination with the calcium chelator BAPTA (10 μM), or the antioxidant N-acetyl-L-cysteine (NAC, 1 mM); scale bar= 10 μm. (B) The graph reports GFP-TFEB nuclear localization, measured as nuclear to cytoplasm fluorescent signal ratio. Data are expressed as mean ± SEM, n=3. (C) We studied GFP-TFEB nuclear localization in HeLa cells silenced for MCOLN-1 or calcineurin-1 upon starvation or 2 hours treatment with sephin1; scale bar= 20 μm. (D) The graph reports GFP-TFEB nuclear localization, measured as nuclear to cytoplasm fluorescent signal ratio. Data are expressed as mean ± SEM, n=4-5. (E) qRT-PCR analysis of the levels of PP3CB and PPR1, respectively the catalytic and the regulatory subunit of calcineurin-1, and MCOLN-1 mRNAs. Data show $2^{-\triangle\triangle Ct}$ and are expressed as mean ± SEM, n =4.

**Supplementary figure 12.** (A-B) We treated HeLa cells for 2 hours with torin-1 (500 nM) or sephin1 (100 μM). We processed the cells upon a 2 hours or 3 hours long wash-out. qRT-PCR analysis of the levels of key TFEB targets right upon 2 (A) and 3 hours long (B) wash-out. Data show $2^{-\triangle\triangle Ct}$ and are expressed as mean ± SEM, n =4. (C) We treated HeLa cells for 2 hours with vehicle or sephin1 (100 μM). We processed the cells upon a 6 hours long wash-out. qRT-PCR analysis of the levels CHOP mRNAs. Data show $2^{-\triangle\triangle Ct}$ and are expressed as mean ± SEM, n =6. (D) We treated HeLa cells for 2 hours with vehicle or cycloheximide (CHX, 100 μg/mL). We processed the cells upon a 6 hours long wash-out. The graph shows the qRT-PCR analysis of the levels of key TFEB targets. Data show $2^{-\triangle\triangle Ct}$ and are expressed as mean ± SEM, n =4-6. (E) We assessed autophagy by monitoring LC3I and LC3B II levels in HeLa cells treated for 2 hours with trehalose (100 mM) or vehicle, followed by 8 hours long wash out and eventually exposed to bafilomycin 1 (baf1, 100nM 2 hours). (F-G) The graphs report the quantification of LC3B II levels, normalized to α-tubulin (F) and LC3II/LC3I ratio (G). Data are expressed as mean ± SEM, n =6.

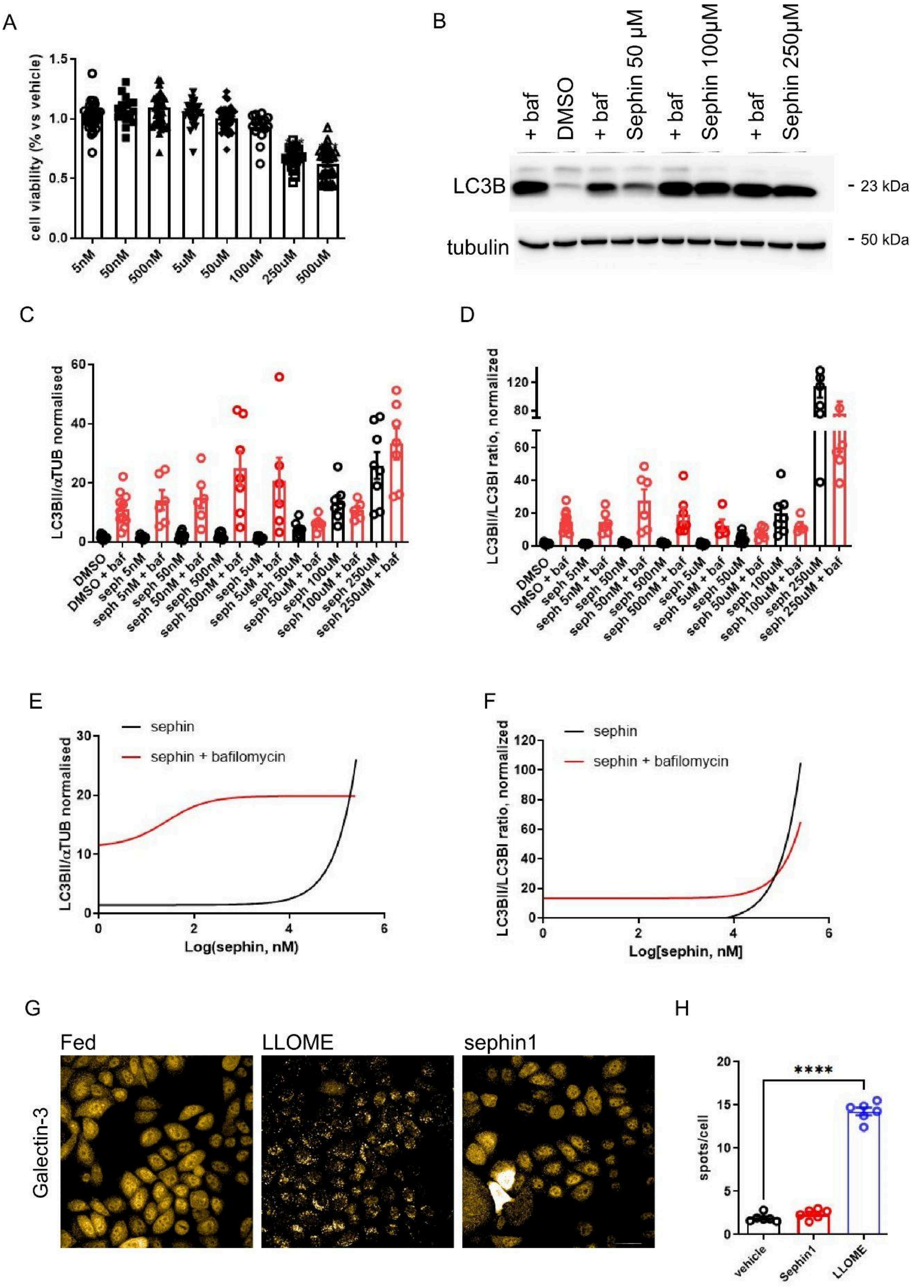


**Supplementary figure 1**

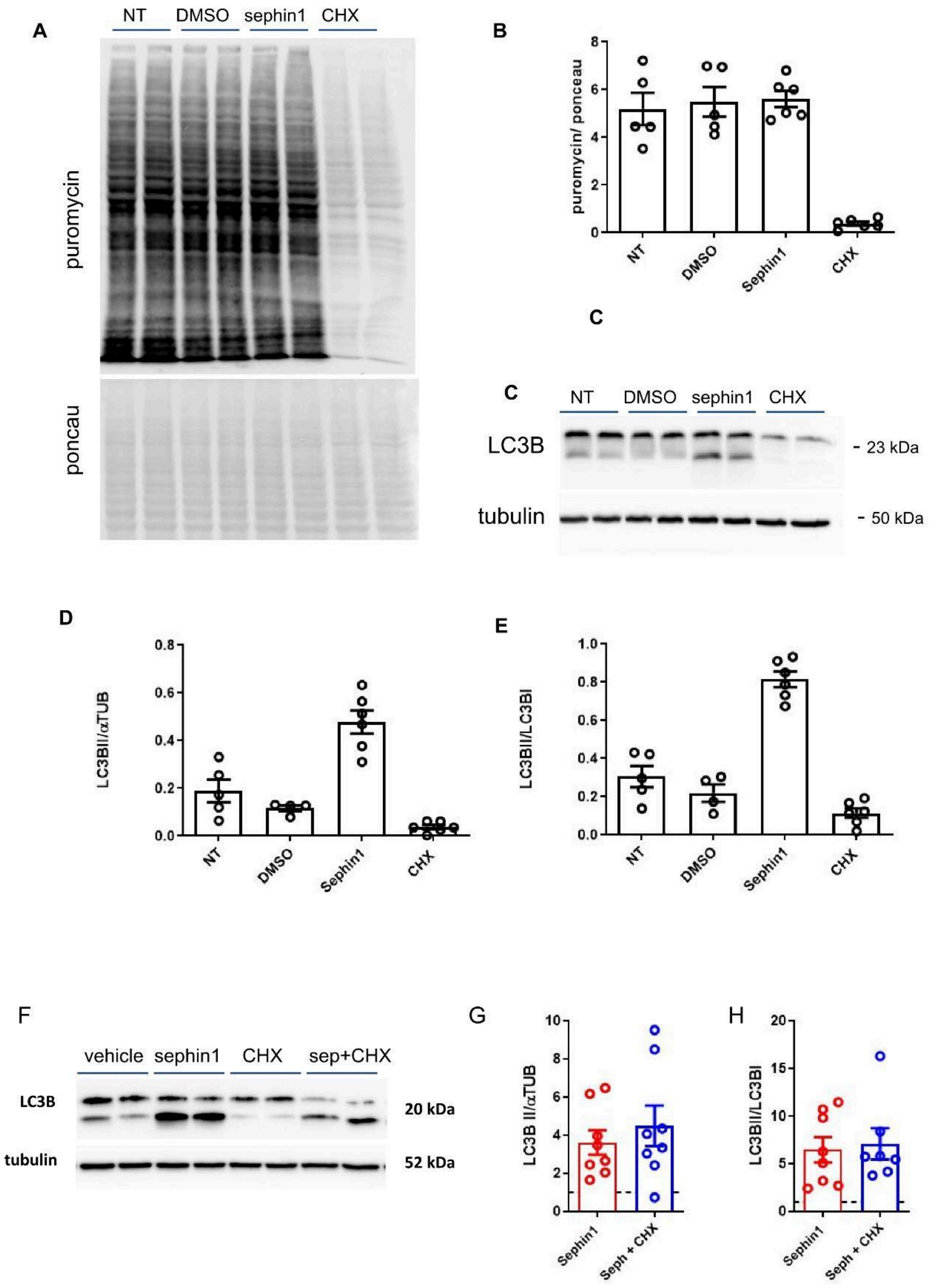


**Supplementary figure 2**

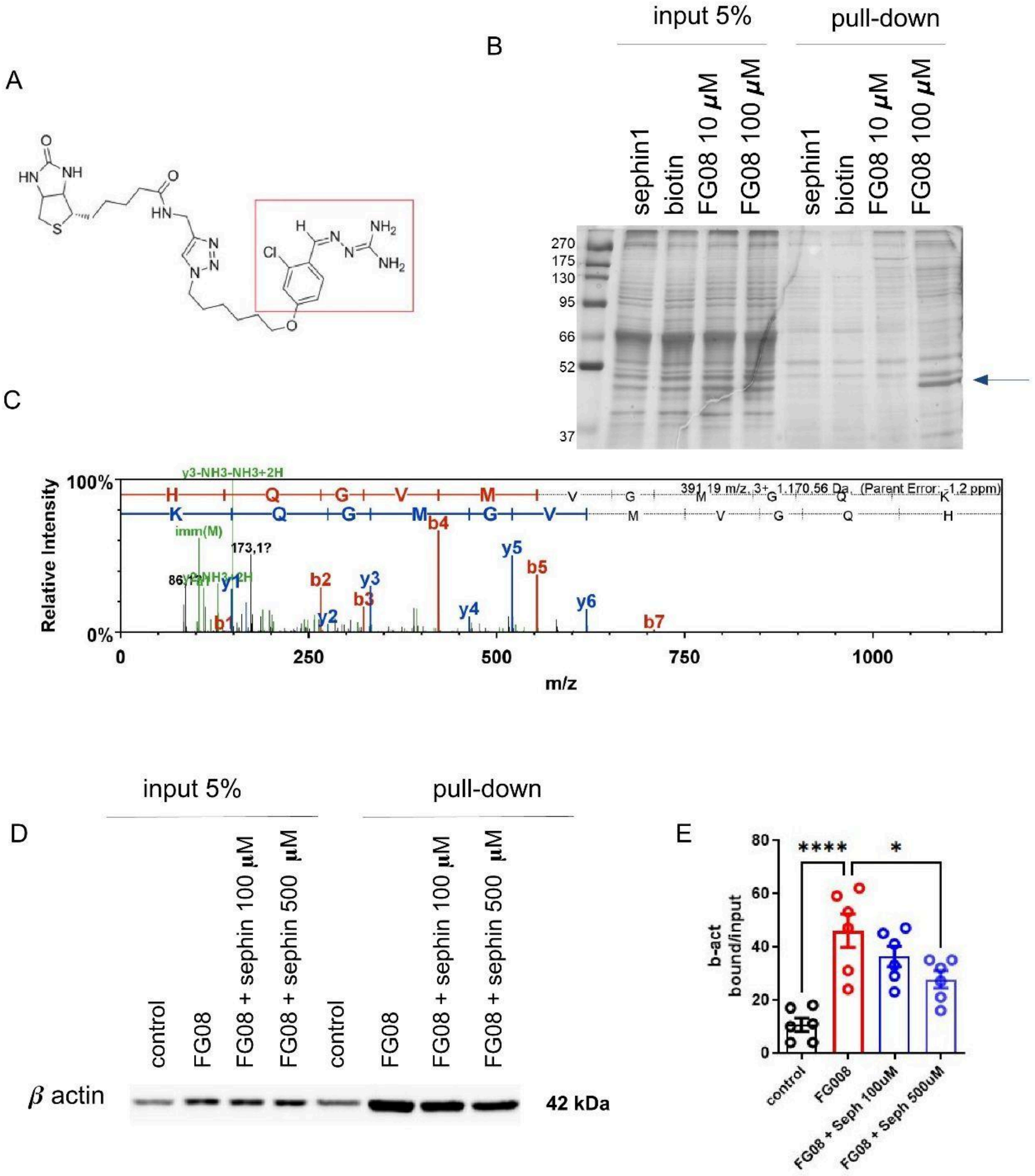


**Supplementary Figure 3**

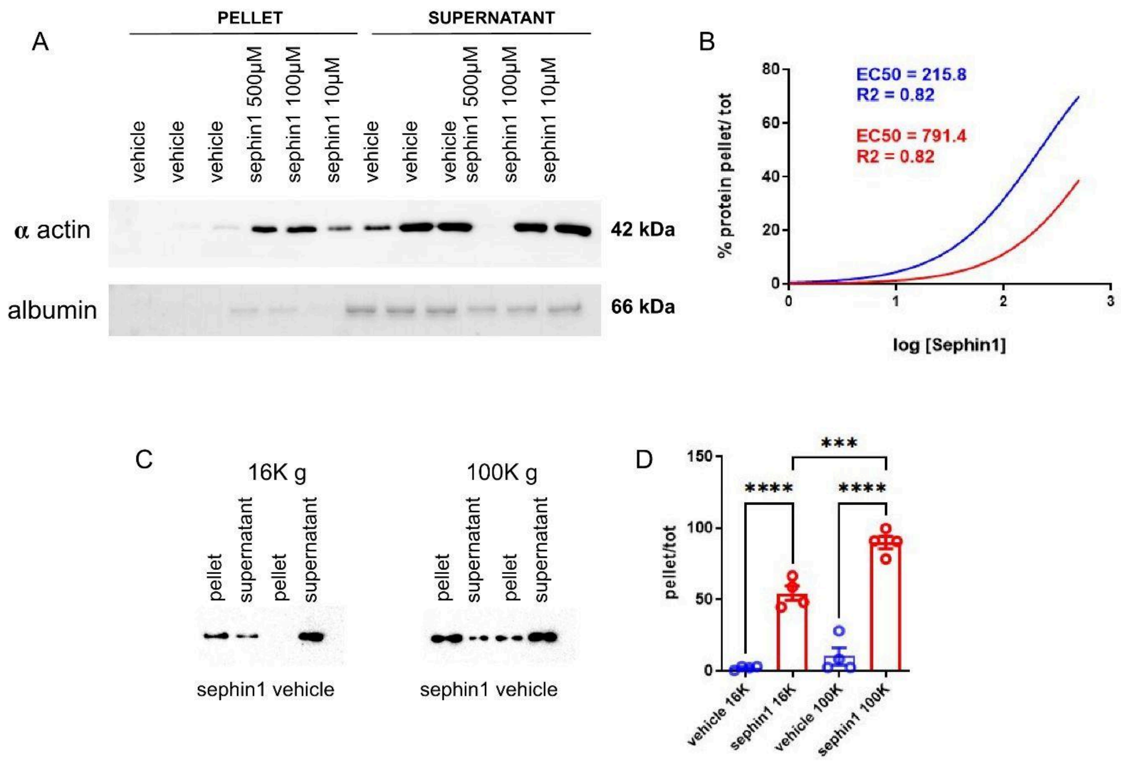


**Supplementary Figure 4**

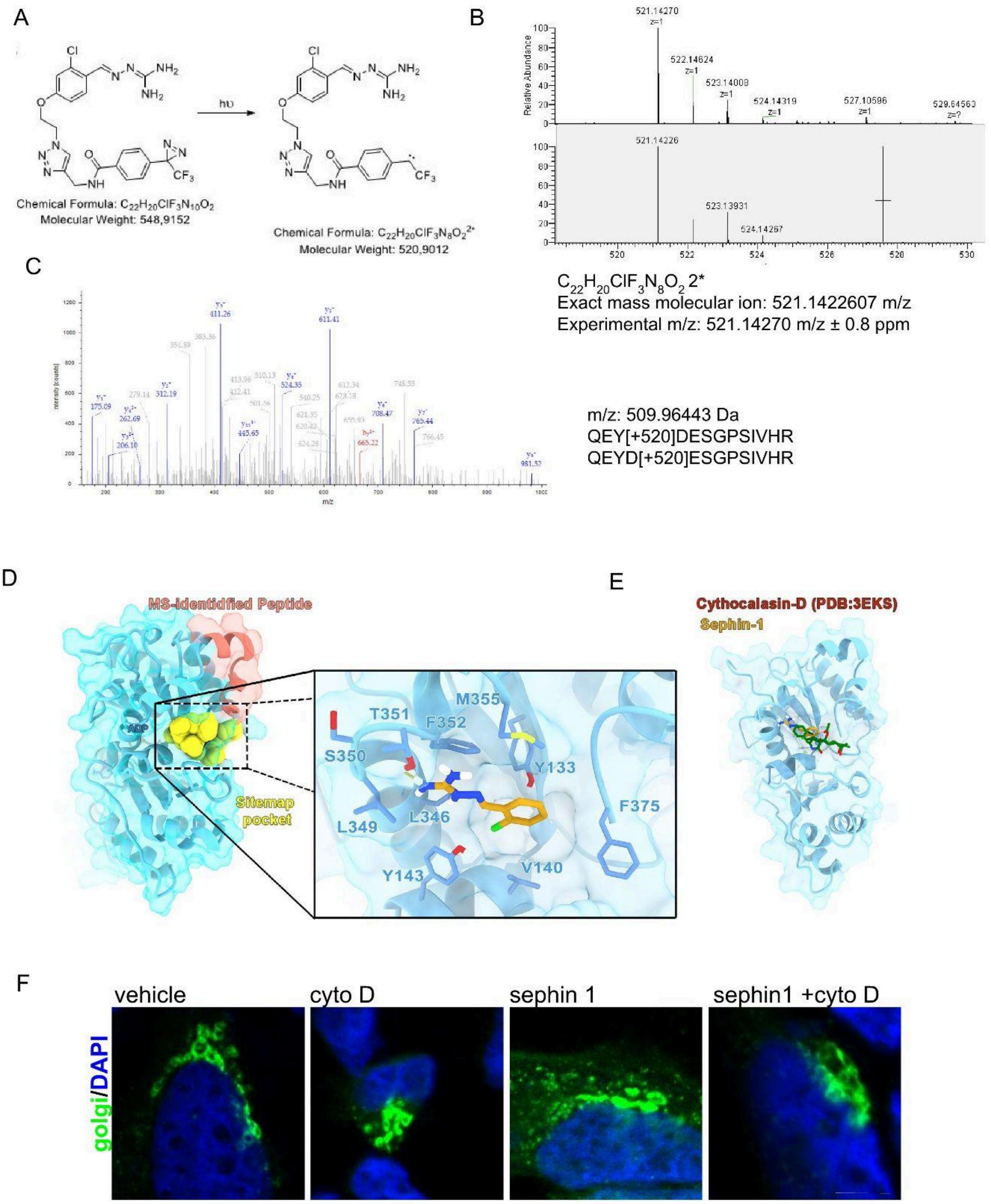


### Supplementary Figure 5

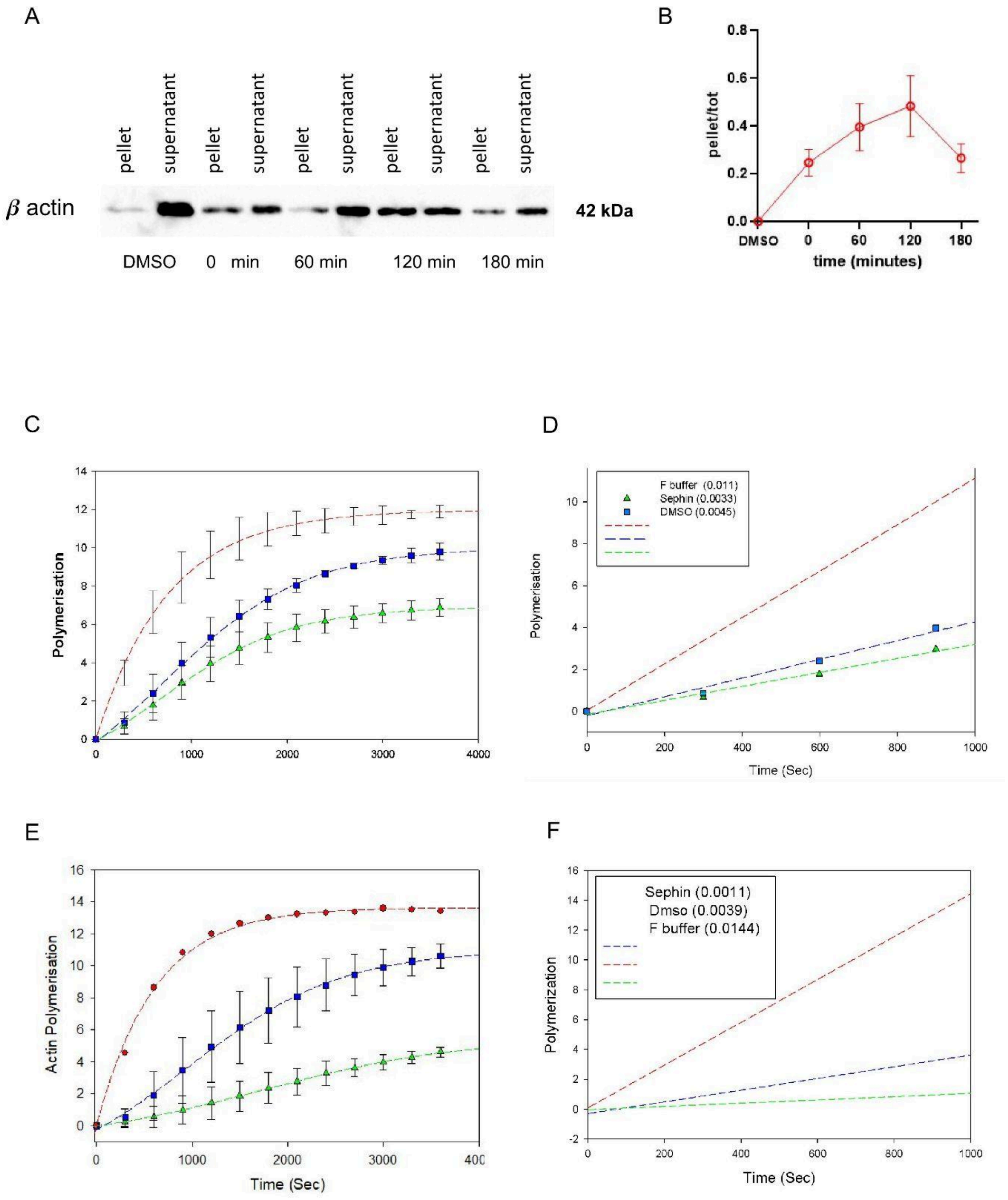


**Supplementary Figure 6**

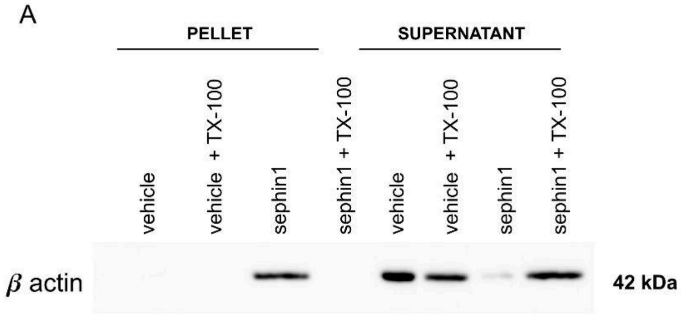


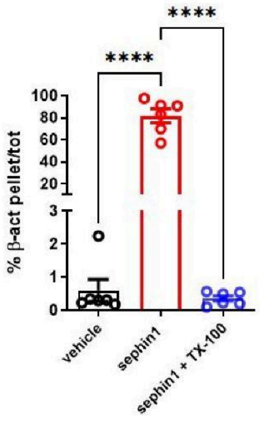


C

| Peptide | Start | End | sephin1 vs DMSO Log10 Fold | sephin1 vs DMSO Pval | sephin1 vs DMSO Adj.Pval |
|---|---|---|---|---|---|
| ALVVDNGSGMCK | 7 | 18 | -1.369124879 | 0.003214601 | 0.033753307 |
| NTVLSGGTTMYPGIADR | 297 | 312 | -1.788469539 | 0.001020283 | 0.021425939 |
| QQMWISK | 353 | 369 | -1.054648068 | 0.030975951 | 0.133794479 |

**Supplementary Figure 7**

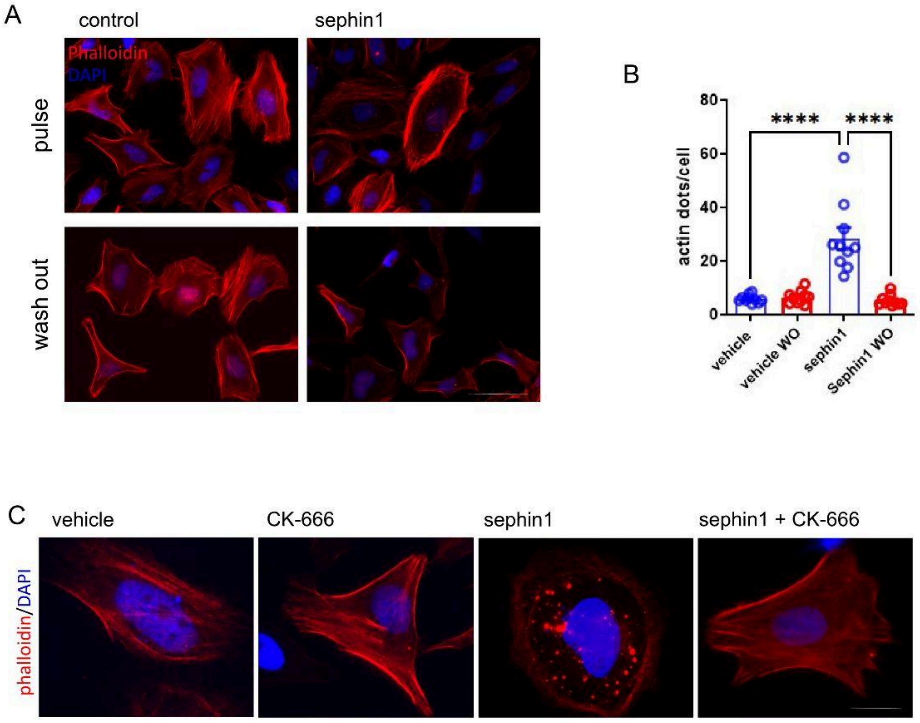


**Supplementary Figure 8**

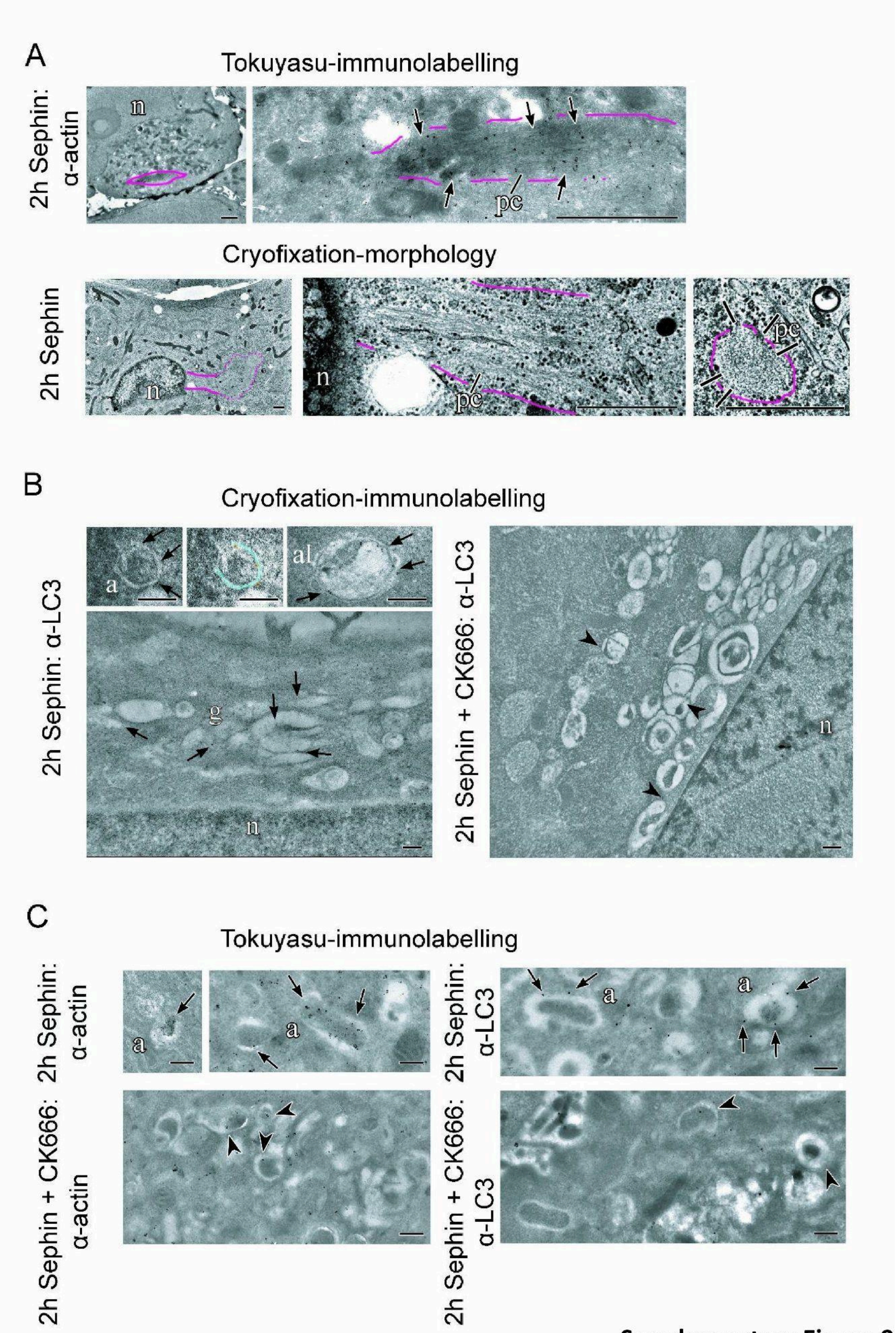


**Supplementary Figure 9**

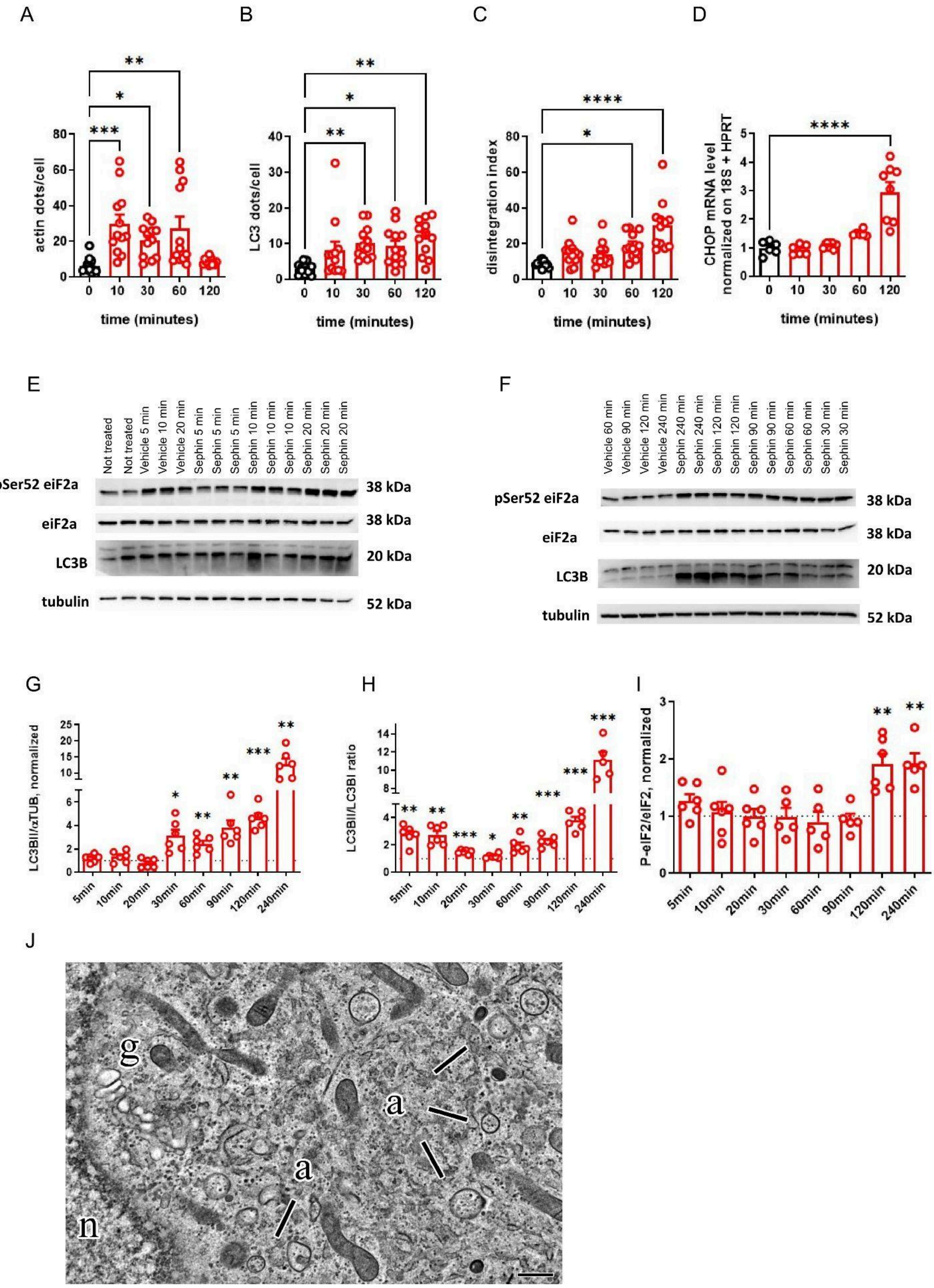


**Supplementary Figure 10**

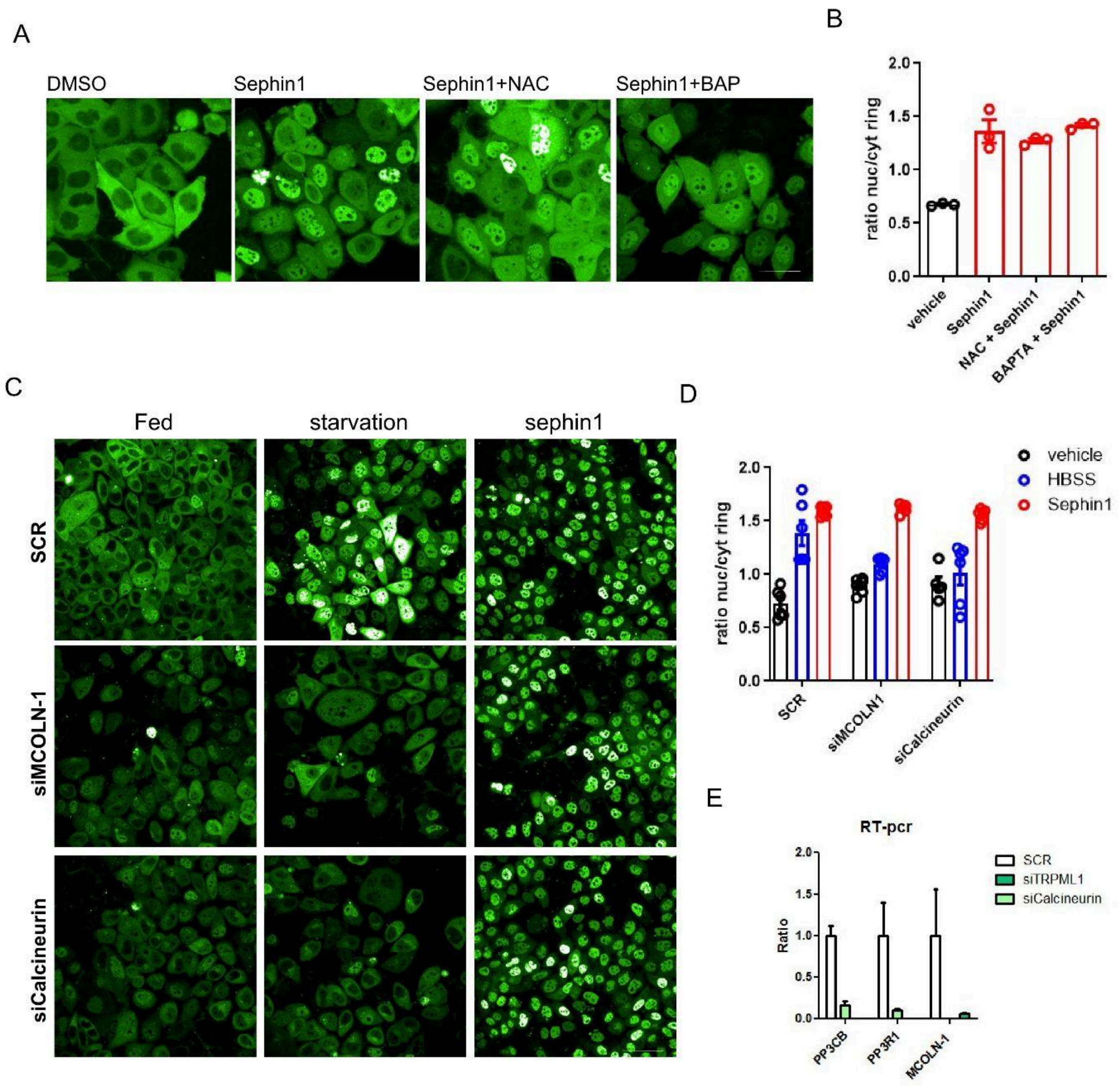


**Supplementary figure 11**

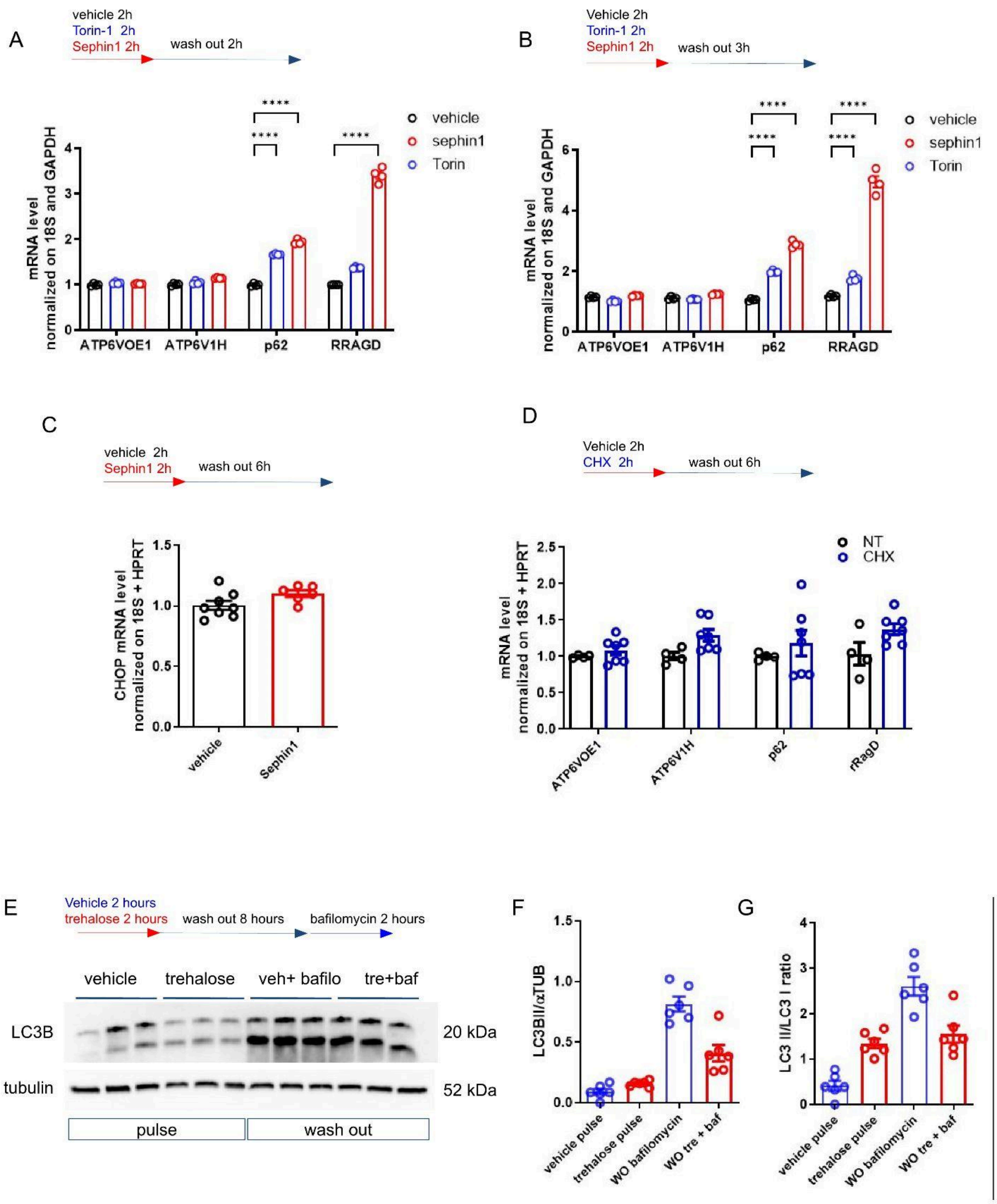


**Supplementary figure 12**
